# The contribution of cohesin to chromatid organisation is critical during chromosome segregation

**DOI:** 10.1101/2021.08.27.457984

**Authors:** Jonay Garcia-Luis, Hélène Bordelet, Agnès Thierry, Romain Koszul, Luis Aragon

## Abstract

Chromosome segregation requires both the separation of sister chromatids and the sustained condensation of chromatids during anaphase. In yeast cells, cohesin is not only required for sister chromatid cohesion but also plays a major role in determining the structure of individual chromatids in metaphase. Separase cleavage is thought to remove all cohesin complexes from chromosomes to initiate anaphase. It is thus not clear how the length and organisation of segregating chromatids are maintained during anaphase in the absence of cohesin. Here we show that degradation of cohesin at the anaphase onset causes aberrant chromatid segregation. Hi-C analysis on segregating chromatids demonstrates that cohesin depletion causes loss of intrachromatid organisation. Surprisingly, TEV-mediated cleavage of cohesin does not dramatically disrupt chromatid organisation in anaphase, explaining why bulk segregation is achieved. In addition, we identified a small pool of cohesin complexes bound to telophase chromosomes in wildtype cells and show that they play a role in the organisation of centromeric regions. Our data demonstrate that in yeast cells, cohesin function is not over in metaphase, but extends to the anaphase period when chromatids are segregating.

**One sentence summary:** Cohesin complexes on yeast chromosomes provide organisation to segregating chromatids.

## Introduction

Mitotic chromosome condensation solves two structural problems during chromosome segregation: firstly, it eliminates entanglements between replicated chromatids, and secondly, it compacts chromosomes so that they avoid being cut by the cytokinetic furrow at the end of mitosis (Guacci et al., 1994). Therefore, it is very important that cells control chromosome length during anaphase, ensuring that chromatids are sufficiently compacted.

Cohesin complexes were first proposed to impact on genome architecture both by providing cohesion between sister chromatids (Guacci et al., 1997; Michaelis et al., 1997) and promoting chromosome condensation during mitosis (Guacci et al., 1997). The discovery that separase cleaves cohesin’s kleisin Mcd1/Rad21/Scc1 subunit (thereafter referred to as Mcd1) (Uhlmann et al., 1999) and the observation that engineered Mcd1 cleavage in metaphase, using TEV (tobacco etch virus) protease expression and TEV recognition sites in Mcd1, triggered bulk separation of the genome (Uhlmann et al., 2000) established the view that yeast cohesin functions solely holding sister chromatids together prior to their separation. However, recent work has shown that the original observation demonstrating a structural function for yeast cohesin in mitosis (Guacci et al., 1997) was indeed correct. Chromosome capture (Hi-C) techniques have revealed that cohesin mediates intrachromosomal loops that are responsible for compacting yeast chromosome arms in metaphase cells, independently of cohesin’s role in sister chromatid cohesion (Lazar-Stefanita et al., 2017; Schalbetter et al., 2017). In contrast to yeast cells, cohesin is not required to organise the arms of metaphase chromosomes in mammalian cells (Sumara et al., 2000). Although cohesin is fully responsible for the organisation of the genome into loops during interphase (Rao et al., 2017), it dissociates from chromosome arms during prophase (Sumara et al., 2000). At metaphase, mammalian cohesin is only retained at centromeres where it provides cohesion (Sumara et al., 2000), while the related SMC complexes, condensin I and II (CI and CII), take over the role of folding mitotic chromosome arms into loops (Gibcus et al., 2018). CII organises large (200- to 400-kb) loops, while CI sub-divides these into smaller loops (80-kb)(Gibcus et al., 2018). During interphase CI is excluded from the nucleus and CII although nuclear, is prevented from acting on chromosomal DNAs by a nuclear protein, Mcph1 (Yamashita et al., 2011).

Yeast cells lack a condensin II complex, and condensin I’s role in chromosome organisation in mitosis is restricted to the ribosomal gene array (Lazar-Stefanita et al., 2017; Schalbetter et al., 2017). Instead yeast condensin is involved in the separation of chromosome arms (Leonard et al., 2015) through a role that promotes decatenation of sister chromatids (Baxter et al., 2011; Sen et al., 2016). Both cohesin and condensin are thought to organise DNA by a process of loop extrusion (Davidson et al., 2019; Ganji et al., 2018; Kim et al., 2019; Kong et al., 2020).

The demonstration that yeast cohesin organises loops on the arms of metaphase chromosomes (Dauban et al., 2020; Garcia-Luis et al., 2019), together with the limited effect observed for condensin on yeast chromosome condensation (Lazar-Stefanita et al., 2017; Schalbetter et al., 2017) and the cleavage of cohesin by separase at the anaphase onset (Uhlmann et al., 2000) present a conundrum currently unexplained; how is the organisation/compaction of chromatids maintained as they are segregating during anaphase?. Here, we have investigated this question. We have studied the organisation of chromosome arms during segregation while we compromised cohesin function using two contrasted experimental approaches; auxin-mediated degradation and TEV-mediated cleavage of Mcd1. We demonstrate that degradation of Mcd1 severely disrupts chromosome structure and consequently the segregation of chromatids in anaphase. In addition, we show that a population of cohesin complexes are present on segregating chromatids during anaphase/telophase and that they are important for centromere organisation. Together, these findings demonstrate that yeast cohesin has a previously unsuspected role necessary for the faithful segregation of chromatids during anaphase.

## Results

### Rapid degradation of cohesin’s kleisin at the anaphase onset causes mitotic catastrophe

To study the role of cohesin on chromosome organisation during anaphase, we searched for an approach to induce the rapid removal of cohesin’s kleisin subunit, Mcd1. To this aim, we used an auxin-inducible degron allele of Mcd1 (*MCD1-AID*) (Nishimura et al., 2009) which allows rapid degradation by poly-ubiquitylation upon exposure to auxin. To evaluate the effect of Mcd1 degradation during chromatid segregation, we blocked cells using transcriptional depletion of Cdc20 (an activator of the anaphase-promoting complex, APC). Under this experimental condition, cells are arrested with chromosomes bipolarly attached to mitotic spindles. Removal of cohesion by artificial cleavage of Mcd1 in Cdc20-depleted cells triggers an anaphase-like state where chromatids segregate to opposite cell poles (Uhlmann et al., 2000).

In cdc20-arrested cells, we observed full degradation of Mcd1-aid 30-60min after addition of auxin (Fig. 1a), demonstrating the rapid and efficient removal of Mcd1 in this experimental setup. Auxin-mediated degradation of Mcd1 led to severe disruption of nuclear segregation (Fig. 1b), with many nuclei appearing to be stuck in anaphase with elongated nuclear masses for extended periods (Fig. 1b-c). To quantify the segregation defects in these cells, we introduced chromosome tags at different genomic locations, and scored the timing and efficiency of their separation. First, we used tags on the arm (*tetO::469*) and telomere (*tetO::558*) regions of chromosome 5, a small chromosome in the yeast genome (Fig. 1c-d). After 180min of auxin addition to cdc20-blocked cultures, we observed that 48% of the cells were still stuck in anaphase (Fig. 1c), 35% of cells showed correct segregation of arm tags and 20% showed misegregation (Fig. 1c). Segregation errors were even higher for telomeric regions (Fig. 1d) and larger chromosomes (Suppl. Fig. 1a-b). These results demonstrate that rapid removal of Mcd1 by degradation causes a catastrophic anaphase-like state with severely impaired separation of chromatids.

**Figure 1.**
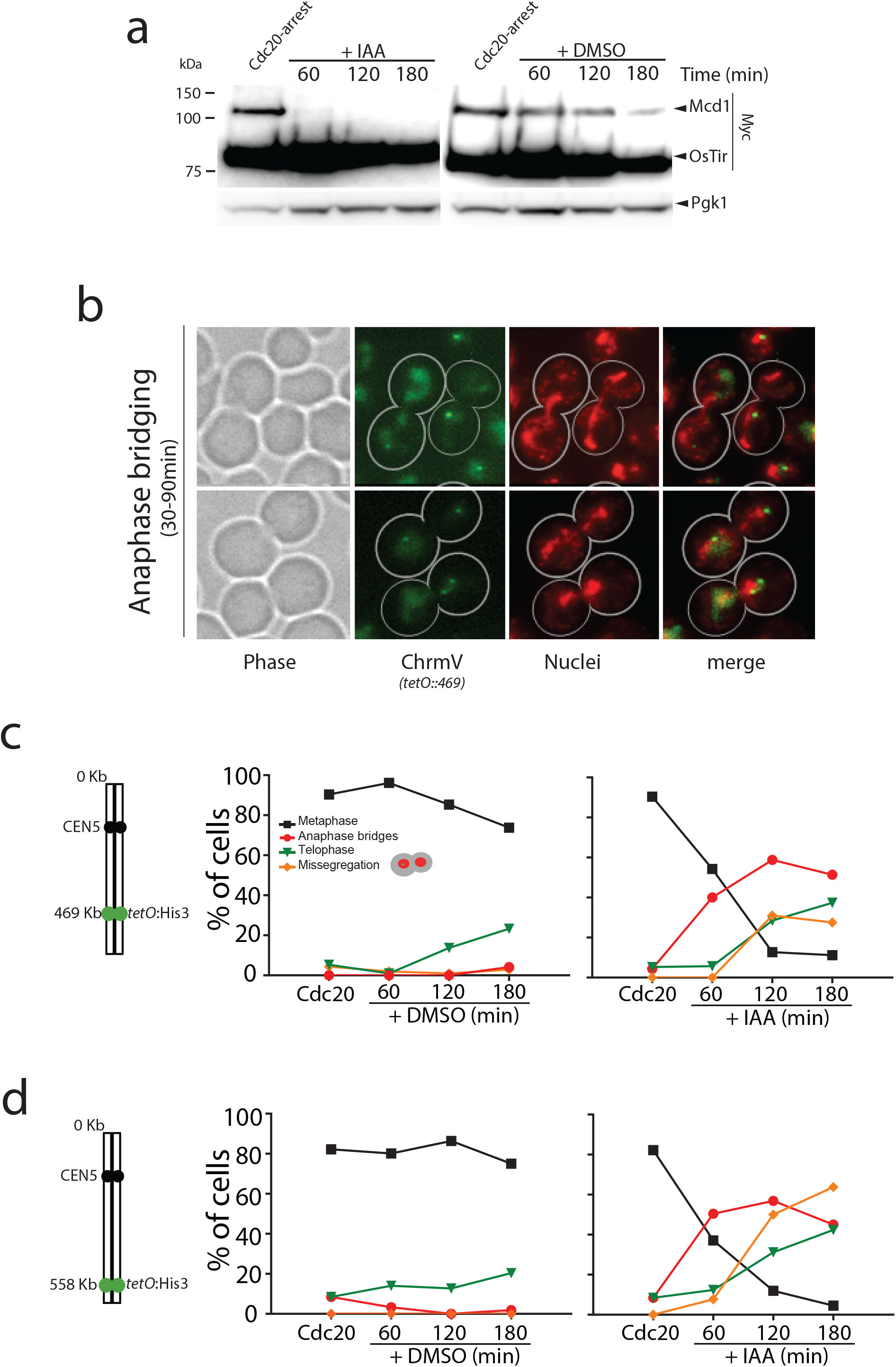
Mcd1 degradation causes catastrophic chromosome segregation. **a,** Cells containing *MCD1* tagged with the auxin degron (*MCD1-AID*) were arrested in metaphase (Cdc20 arrest). The culture was split in two, one half was treated with DMSO 1% and the other with 6mM auxin (IAA) to degrade Mcd1. Samples were taken for and anti-Myc immunoblotting to detect Mcd1. **b,** Representative images of cells 30-90 minutes after degradation of Mcd1. Cells were analysed for nuclear separation (DAPI stain) and chromosome segregation (GFP dots marking the middle of chromosome V; *tet:469*). **c,** Analysis of nuclear and chromosome segregation using DAPI and GFP dots marking the middle of chromosome V (*tet:469*). Experimental conditions for the timecourse are as described in *a*. **d,** Analysis of nuclear and chromosome segregation using DAPI and GFP dots marking the telomeric region of chromosome V (*tet:558*). Experimental conditions for the time-course are as described in *a*. Each time point represents the percentage of cells at the indicated cell cycle stage. At least 100 cells were quantified for each time-point.

### TEV-induced cleavage of cohesin allows bulk genome separation with minor segregation errors

Previous studies have shown that engineered cleavage of Mcd1 in cdc20-arrests, using TEV (tobacco etch virus) protease expression and TEV recognition sites on Mcd1, triggers an anaphase-like division where nuclear masses separate (Uhlmann et al., 2000). We evaluated segregation in TEV-induced anaphases using the previously published protocol (Uhlmann et al., 2000). Induction of TEV expression led to cleavage of Mcd1 after 60 min (Fig. 2a) as expected. Bulk nuclear segregation occurred 90-120 min following the induction (Fig 2b) as it had been previously reported (Uhlmann et al., 2000). Importantly, chromosome tags located on the centromere region of chromosome 5 segregated efficiently (Fig. 2c). Therefore, unlike auxin-mediated degradation of Mcd1, TEV-cleavage allows nuclear segregation.

**Figure 2.**
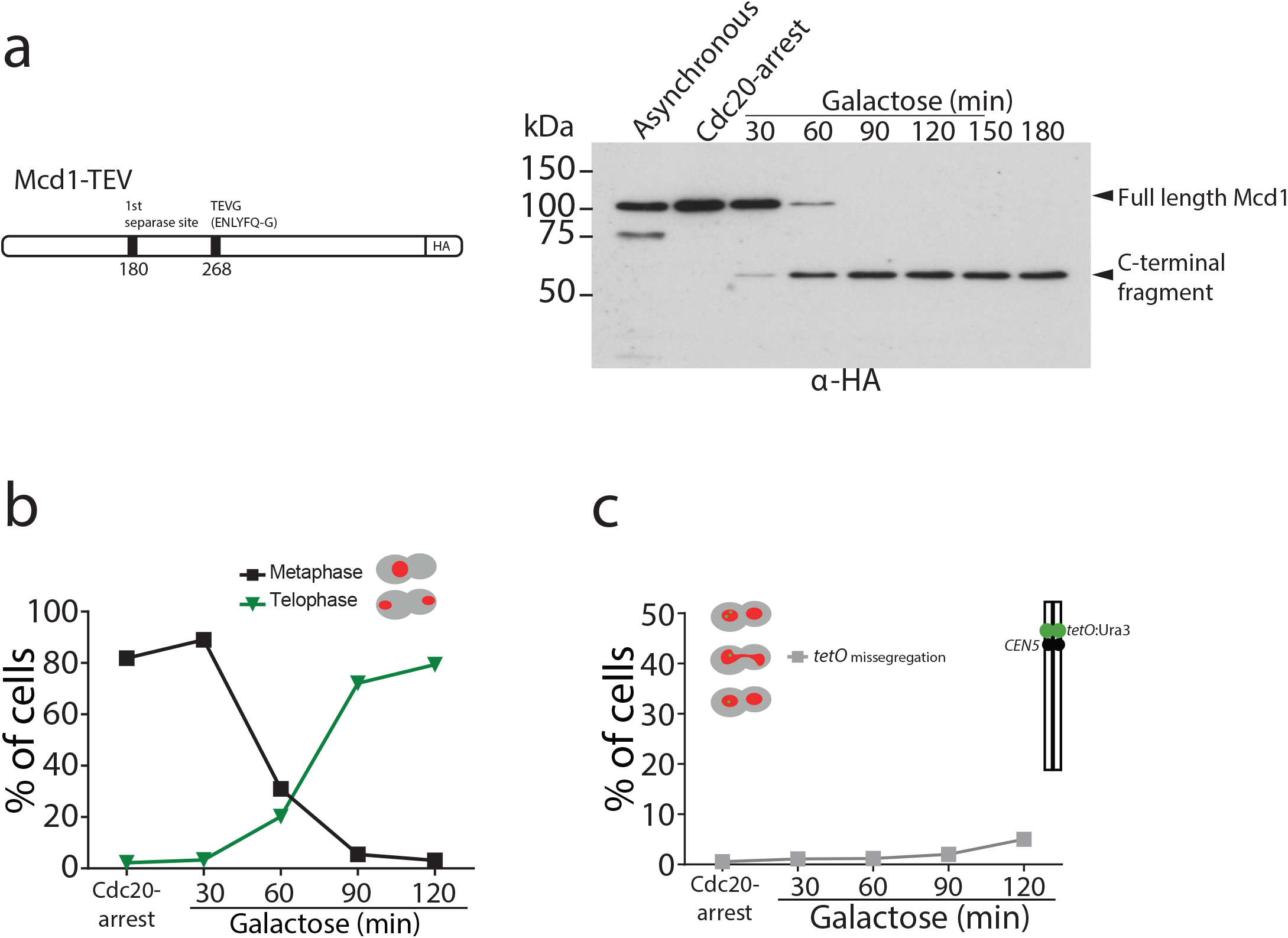
Mcd1 cleavage by TEV protease leads to bulk chromosome segregation. **a**, Schematic of engineered *MCD1* with the wild-type separase cleavage site maintained in the position 180 and the separase cleavage position 268 replaced by the TEV cleavage sequence ENLYFQG (upper panel). Cells were arrested in metaphase (Cdc20 arrest) and induced to express TEV protease. Samples were taken every 30 minutes for 3 hours for immunoblotting against HA epitope to follow the C-terminus of Mcd1. **b**, Nuclear segregation in TEV-induced anaphase. Nuclear segregation was monitored using DAPI staining of cells. Experimental protocol as in *a*. **c**, Chromosome segregation in TEV-induced anaphase. Chromosome tags inserted in a centromere proximal site (Ura3::*tetO*) on chromosome V were monitored for segregation. Experimental protocol as in *a*. Each time point represents the percentage of cells at the indicated cell cycle stage. At least 100 cells were quantified for each time-point

We noticed that in TEV-induced anaphases the cleaved C-terminal fragment of Mcd1 was fully stable during the entire time-course (Fig. 2a). The lack of Mcd1 fragment degradation after TEV cleavage stems from the fact that TEV protease cleavage occurs following the glutamine (Q) residue of the TEV recognition site (‘ENLYFQ*G’) leaving a glycine (G) amino-acid residue at the N-termini (referred to as TEVG), which is not well recognized by the N-rule pathway (Varshavsky, 2011). In contrast, separase cleavage leaves an arginine (R) residue at the N-termini of the cleaved product (‘SVEQGR*R’), which is a good substrate for N-end rule degradation (Varshavsky, 2011). Interestingly, Beckouet *et al*. have shown that following TEV-induced cleavage not only the C-terminal fragment of Mcd1 is stabilised but also the N-terminal (Beckouet et al., 2016). This raises the possibility that Mcd1 fragments could remain associated to the Smc core subunits following TEV cleavage. To test whether this is the case, we tagged Smc3 with the V5 epitope and Mcd1 with FLAG and HA tags at the N- and C-terminus respectively (Fig. 3). We performed IPs on synchronised TEV-anaphases (Fig. 3) to follow whether the cleaved fragments stay associated to the cohesin Smc core. Both N- and C-terminal Mcd1 fragments were immunoprecipitated by Smc3 after TEV cleavage (Fig. 3 and Suppl. Fig. 2). Therefore, the structural integrity of cohesin tripartite complex remains intact after TEV cleavage of its kleisin subunit. Next, we sought to test whether inducing full degradation of Mcd1 fragments after TEV cleavage generated a similar phenotype to that observed in anaphases induced by degradation of Mcd1-aid with auxin (Fig. 1c).

**Figure 3.**
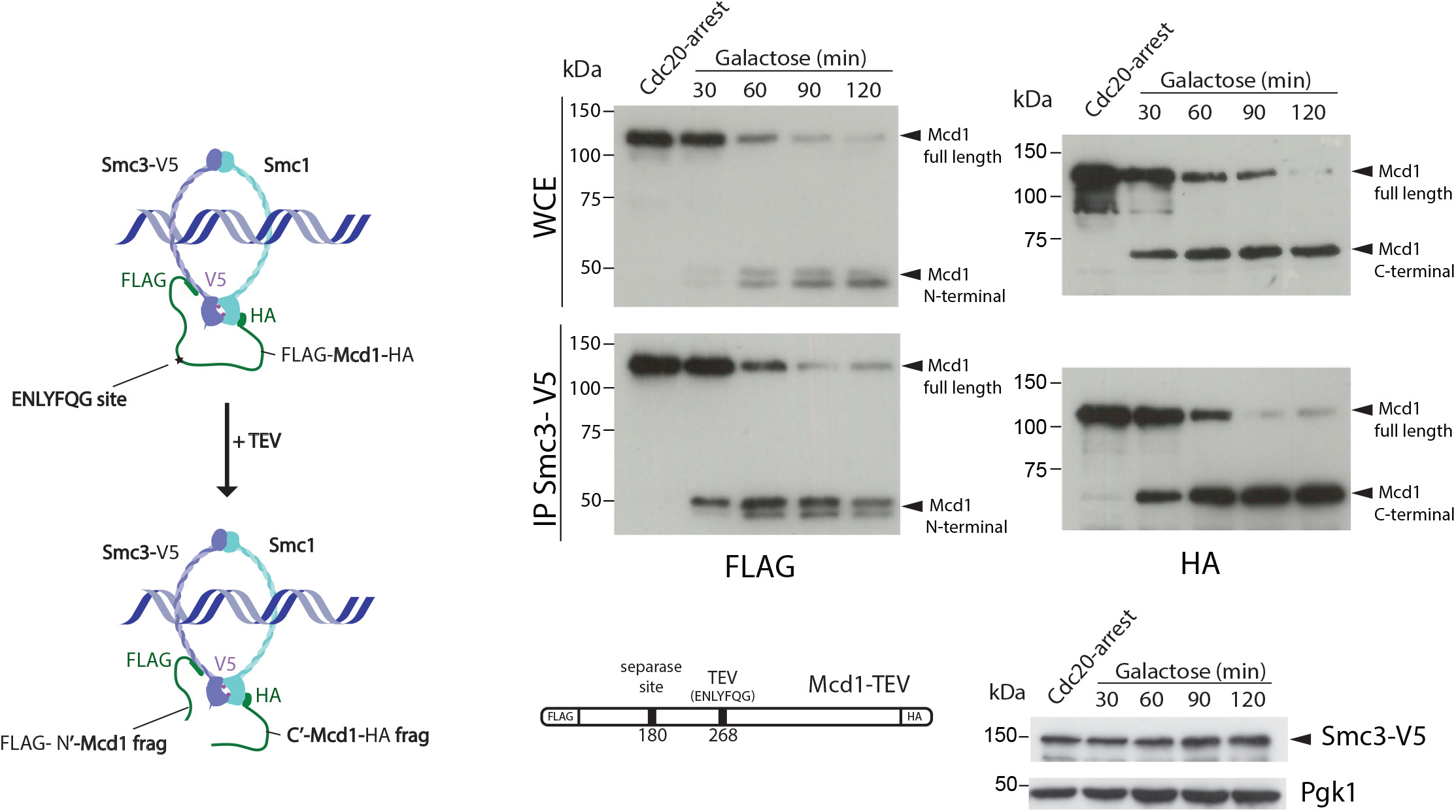
Cohesin ring structure remains after TEV cleavage of Mcd1. Schematic of engineered cohesin with *MCD1* tagged in N-terminus with FLAG, the C-terminus with HA, and with a substitution of the 268 separase cleavage site with the TEV cleavage sequence ENLYFQG. Cells also harboured a copy of Smc3 tagged in C-terminus with V5 (left panel). Cells were synchronized using a Cdc20 arrest and TEV was induced to cleave Mcd1. Samples were taken every 30 minutes for two hours and Smc3 immunoprecipitated using anti-V5 antibody. We used immunoblotting with anti-FLAG and HA antibodies to detect the cleaved fragments of Mcd1 (right panel). Two biological replicas were performed.

To this aim, we used a TEV recognition site on Mcd1 that was able to yield a C-terminal fragment with a terminal amino-acid recognized by the N-end rule pathway. We found this to be the case when we used the TEV recognition site ‘ENLYFQF’ (referred to as TEVF) (Fig. 4a). This site leaves a phenylalanine, rather than a glycine, as the N-terminal aminoacid after TEV cleavage. Importantly, N-terminal phenylalanine is a good substrate for N-end rule degradation (Varshavsky, 2011). We used FLAG and HA tags at the N- and C-terminus of Mcd1, respectively, to detect the two products generated by TEV cleavage and compared their stability in Mcd1 proteins containing the classical ‘ENLYFQG’ (TEVG) or ‘ENLYFQG’ (TEVF) sites (Fig. 4a-b). Cleavage of TEVG generated stable N- and C-terminal fragments observed before (Fig. 4b). In contrast, both N- and C-terminal fragments were rapidly degraded after cleavage on TEVF sites (Fig. 4a).

**Figure 4.**
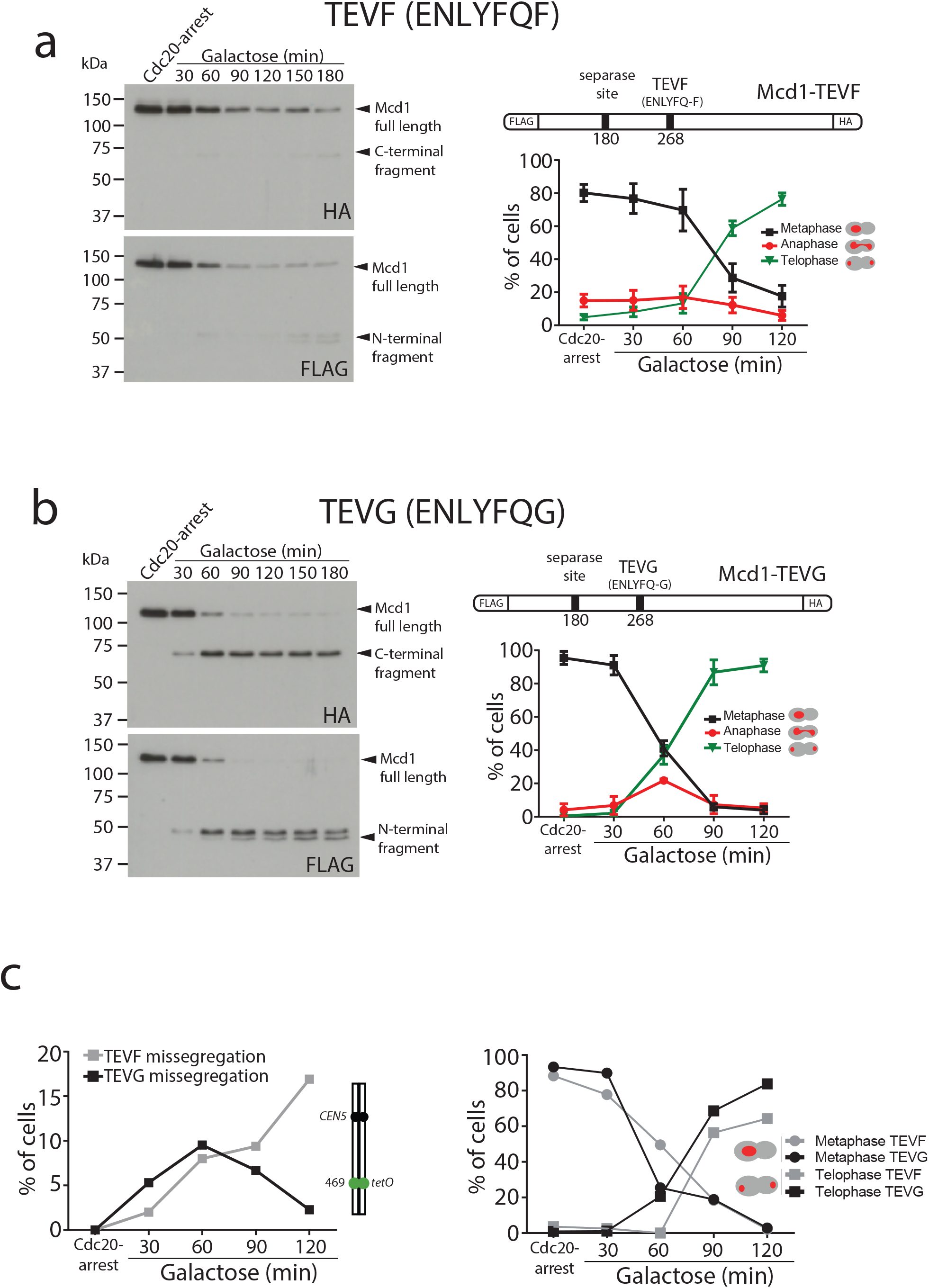
Degradation of Mcd1 fragments after TEV cleavage affects segregation efficiency. **a**, Cells with *MCD1* tagged at its N-terminus with FLAG and its C-terminus with HA, and with the 268 separase cleavage site replaced by the TEV recognition site ENLYFQF (TEVF) (top right diagram) were arrested in Cdc20 before TEV induction. Samples were taken every 30 minutes for two hours and immunoblotted against FLAG or HA to detect Mcd1 N-terminus and C-terminus cleaved fragments respectively (left). Nuclear segregation was monitored during the TEV-induced anaphase with DAPI staining (bottom right graph). **b**, Cells with *MCD1* tagged at its N-terminus with FLAG and its C-terminus with HA, and with the 268 separase cleavage site replaced by the TEV recognition site ENLYFQG (TEVG) (top right diagram) were arrested in Cdc20 before TEV induction. Samples were taken every 30 minutes for two hours and immunoblotted against FLAG or HA to detect Mcd1 N-terminus and C-terminus cleaved fragments respectively (left). Nuclear segregation was monitored during the TEV-induced anaphase with DAPI staining (bottom right graph). **c**, Cells carrying *MCD1* with the 268 separase cleavage site substituted for either TEV cleavage ENLYFQF (TEVF) or ENLYFQG (TEVG) were treated as in *a* and *b* and monitored for nuclear and chromosome segregation using DAPI and GFP dots marking the middle of chromosome V (*tet:469*). Each time point represents the percentage of cells at the indicated cell cycle stage. Error bars represent the SD of 3 biological replicas. At least 100 cells were quantified for each time-point

Next, we compared segregation kinetics in TEV-induced anaphases with TEVG and TEVF cleavage sites. Bulk nuclear separation was observed in both conditions (Fig. 4a-b), with most cells showing full segregation after 120 min of TEV induction (>85% in TEVG and >75% in TEVF). However, we noticed minor delays during anaphase progression in cells with TEVF recognition sites (Fig. 4a). Next, we compared the fidelity of segregation in TEVG and TEVF anaphases using tetO:469kb tags inserted in the middle of chromosome 5. As observed previously, cells carrying TEVG recognition sites segregated tags correctly (with <5% missegregation observed) (Fig. 4c). In contrast, tag missegregation was observed in 16% of telophases when TEVF recognition sites were present on Mcd1 (Fig. 4c and Suppl. Fig. 3). These results demonstrate that degradation of Mcd1 fragments after TEV cleavage affects the fidelity of chromosome segregation but, unlike Mcd1 degradation, does not severely prevent bulk nuclear separation.

### Depletion or cleavage of cohesin differentially affect chromatin structure and segregation

Cohesin mediates intrachromosomal loops in metaphase arrested chromosomes, providing a structural framework for the compaction of chromosome arms (Lazar-Stefanita et al., 2017; Schalbetter et al., 2017). How chromosomes are organised during yeast anaphase, when individual chromatids are being pulled to the poles, is not well understood. Since the cohesin tripartite complex remains associated after TEV cleavage (Fig. 3), we considered the possibility that TEV-cleaved cohesin retains a role in maintaining the structure of segregating chromatids. To further investigate this, we first performed ChIP analysis of Smc3 during TEV-induced anaphases along chromosome 5 (Fig. 5a). We observed that significant levels of chromatin-bound Smc3 remained during initial timepoints (Fig. 5a; TEV-induced). In contrast, Smc3 was rapidly lost from chromatin in anaphases induced by auxin-mediated degradation of Mcd1 (Fig. 5a; auxin-induced). Having observed that TEV-cleaved cohesin remains chromatin-bound for significantly longer periods than auxin-degraded cohesin (Fig. 5a), we decided to use these two conditions to investigate the organisation of chromatid structure. To this aim, we built Hi-C libraries from timepoints when the bulk of Mcd1 had been either TEV-cleaved or auxin-degraded (Suppl. Fig. 4). Following sequencing, we computed the corresponding normalized genome-wide contact maps for Mcd1-TEV and Mcd1-aid (Fig. 5b-c). When we compared the contact maps of individual chromosomes obtained from cells arrested in metaphase using Cdc20 depletion (Fig. 5b), to those obtained for Mcd1-TEVG and Mcd1-AID during the induced anaphases, we observed a decrease in intrachromosomal contacts (Fig. 5b-c, Suppl. Fig. 5). The contact probability (P) as a function of genomic distances of all chromosome arms showed a reduction of contacts in the 10-30kb range for Mcd1-TEVG and Mcd1-AID samples compared to Cdc20 arrests (Fig. 5c, Suppl. Fig. 5). Notably, the reduction in Mcd1-AID was significantly more pronounced than in Mcd1-TEVG (Fig. 5c, Suppl. Fig. 5). Therefore, these results indicate that a structural role persists after TEV cleavage, but not Mcd1 degradation.

**Figure 5.**
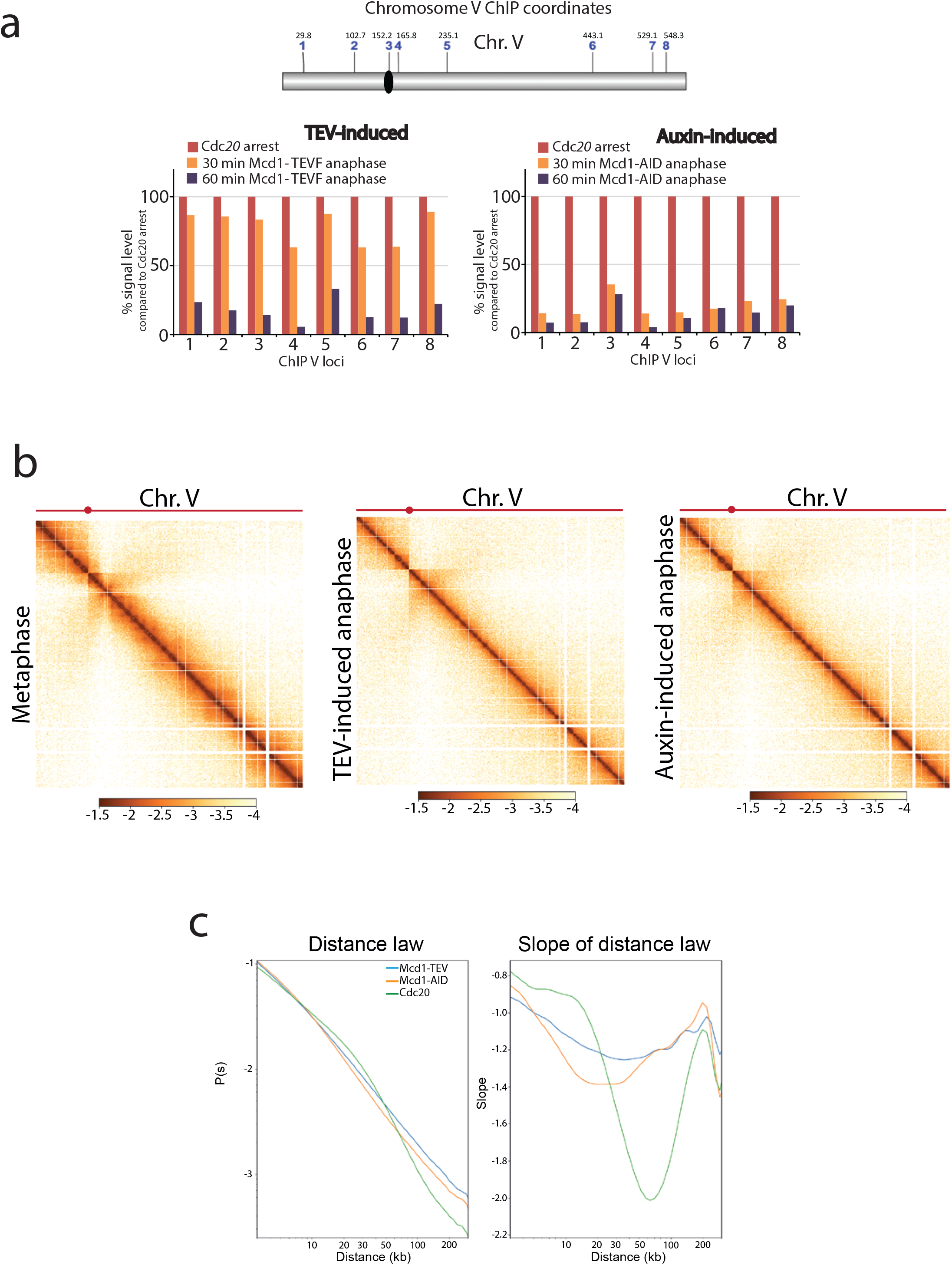
Cohesin presence on segregating chromatids is necessary for *cis* interactions. **a**, Chromatin-immunoprecipitation analysis (ChIP) of Smc3-V5 binding along chromosome V of cells arrested in metaphase (Cdc20 arrest) containing either *MCD1* with the 268 separase cleavage site substituted by the TEV recognition sequence ENLYFQF (TEVF) or with *MCD1-AID*. Samples were taken every 30 minutes for one hour after induction of the TEV protease or addition of the auxin IAA respectively and analysed. Each bar represents the average of 2 technical replicas. **b**, Cells containing either *MCD1-TEVG or MCD1-AID* were arrested in Cdc20 metaphase arrest and MCD1 was cleaved or degraded respectively. Samples for HiC analysis were taken (*MCD1 TEVG* 90 min; *MCD1-AID* 60 minutes). Contact maps (bin = 1kb) of chromosome V from cell populations are shown. Brown to yellow colour scales represent high to low contact frequencies, respectively (log10). A Cdc20 metaphase arrest was also processed as a reference **c**, Average intra-chromosomal contact frequency (P) between two loci with respect to their genomic distance (s) along the chromosome of cell populations treated as in “a” (left). Derivative of P(s) curve (right). Each condition has 2 biological replicas.

### Cohesin organises centromeres of telophase arrested chromosomes

Our results demonstrate that the fidelity of chromosome segregation requires the maintenance of cohesin-dependent structure during anaphase. Next, we investigated whether cohesin complexes are removed in fully segregated chromosomes when they reach the cell poles in telophase. To this aim, we first fused the green fluorescent protein (GFP) to the cohesin subunit Smc3 and imaged its localisation on telophase arrested cells (Fig. 6a). Smc3-GFP signal was observed on segregated nuclei, with a discrete dot present at the cell poles that colocalised with the spindle pole body protein Spc29 (Spc29-RedStar2) (Fig. 6a). This suggests that Smc3-GFP might be enriched at centromeric regions of telophase chromosomes. To confirm this possibility, we used calibrated ChIP-seq in *cdc15-2* arrested cells (Suppl. Fig. 6) to identify whether centromeric regions and any other potential genomic sites are bound by cohesin. To have a guide for the positions where cohesin is normally enriched, we used Smc1 localisation on cells arrested in metaphase where cohesin binding is maximal (Fig. 6b). To ensure that signals detected in telophase arrests were not due to background noise, we subtracted the signal obtained using untagged cells in our analysis (Fig. 6b). The number of cohesin binding sites per chromosome was dramatically reduced in telophase arrested cells (Fig. 6b), and only a few arm sites exhibited levels above background (Fig. 6b). Comparison of average profiles across CEN sites confirmed our cytological results demonstrating that cohesin is indeed enriched at these regions in telophase arrested chromosomes (Fig. 6c, Suppl. Fig. 7).

**Figure 6.**
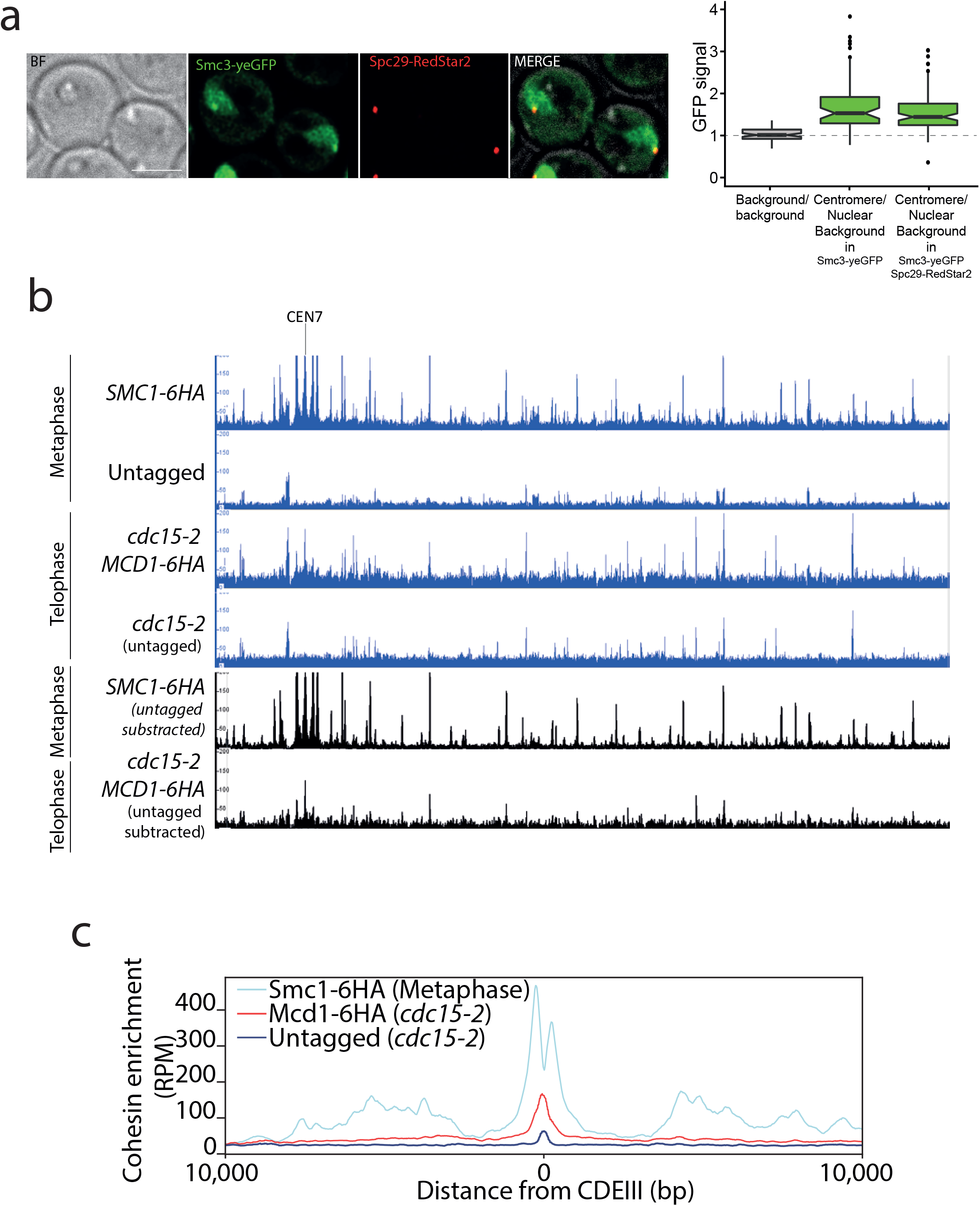
Cohesin is present around centromere regions in telophase arrested cells. **a**, Cells containing *CDC15-AID* and the tagged cohesin subunit *SMC3-yeGFP* were arrested late anaphase and the GFP signal at the centromere was calculated as a ratio comparing it to the nuclear background signal. Cells also carried Spc29-RedStar2, a spindle pole body component that was used as spatial reference to determine colocalization with centromeres. At least two biological replicas were done for each condition. At least 20 cells were quantified in each replica. **b**, Enrichment of cohesin along *S. cerevisiae* chromosome 7 measured by calibrated ChIP-seq in cells containing Smc1-6HA arrested in metaphase (nocodazole), untagged cells arrested in metaphase (nocodazole), cells containing *MCD1-6HA* arrested in late anaphase (*cdc15-2*) and untagged cells arrested in telophase (*cdc15-2*; top 4 blue lanes). The two bottom black lanes show the enrichment of cohesin subunit Smc1-6HA and Mcd1-6HA after subtraction of ChIP-seq signal of the untagged cells arrested in metaphase or telophase (*cdc15-2*). *CEN7* marks the location of the centromere. ChIP-seq for each condition has two biological replicas **c**, Average calibrated ChIP-seq profiles of Smc1-6HA (metaphase arrest) and Mcd1-6HA (telophase arrest, *cdc15-2*) from the centromere *CDEIII* region of the 16 chromosomes is shown.

Next, we sought to test whether centromere-bound cohesin contributes to the organisation of these regions in telophase. We arrested cells using an analogue-sensitive (AS) allele of Cdc15, and inactivated cohesin using the Mcd1-AID and Smc3-AID alleles after telophase arrest had been achieved (Fig. 7a). We then build Hi-C libraries from cells arrested in telophase with and without degradation of cohesin after the arrest (Suppl. Fig. 8). Comparison of the contact maps revealed changes at centromeric regions in telophase cells depleted for cohesin (Fig. 7b), including a decrease in isolation of *CEN*-proximal regions from other sites down chromosome arms *in cis* (Fig. 7c). These results demonstrate that cohesin complexes influence intrachromatid contacts at centromeres in telophase chromosomes. Moreover, the *in trans* interaction of *CEN* sequences was also reduced in *cdc15-as* cells with depleted cohesin (Fig. 7d). Therefore, inactivating cohesin in telophase also reduces centromere clustering of the yeast genome.

**Figure 7.**
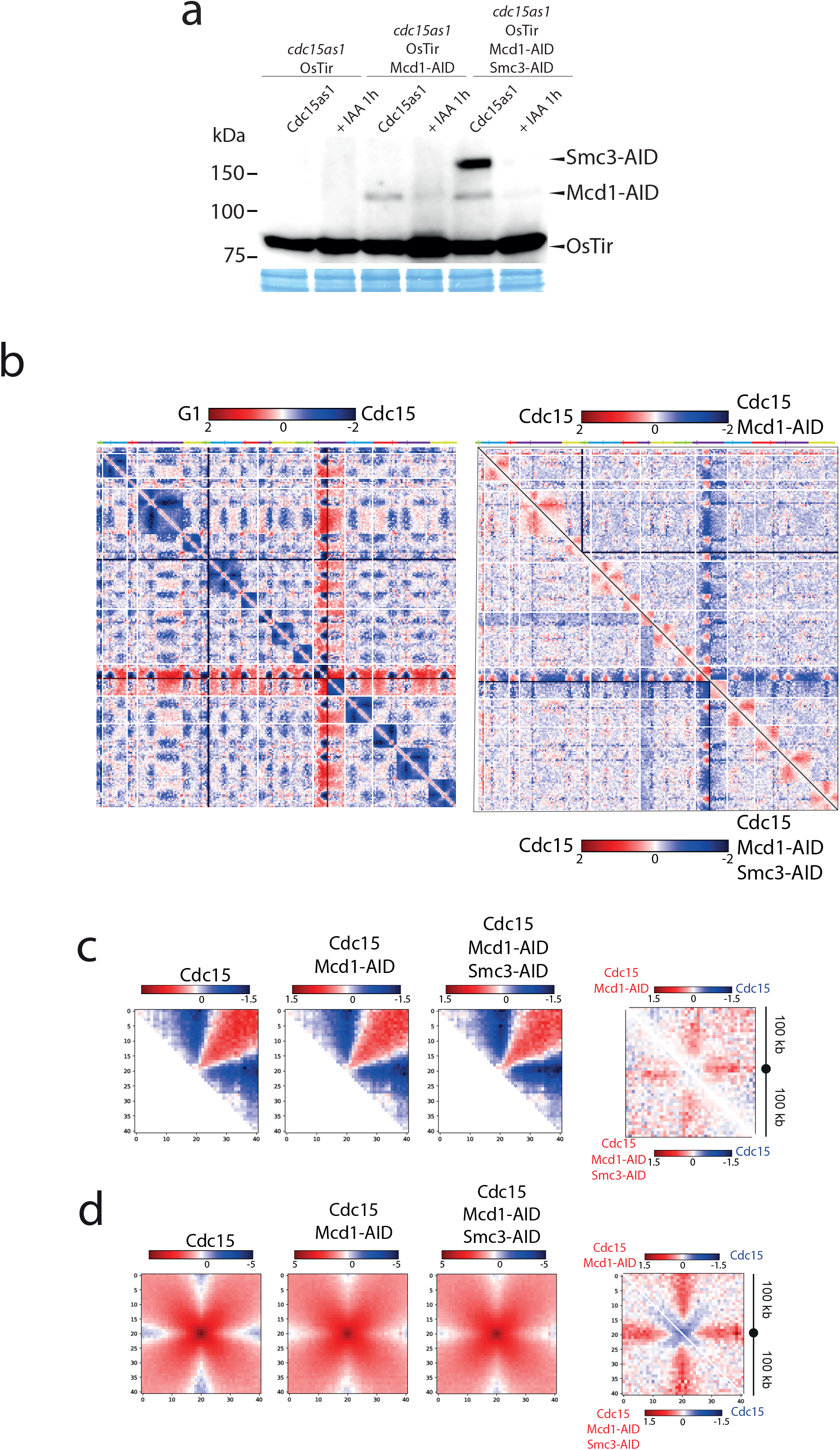
Cohesin organises centromere regions in telophase arrested cells. **a**, Degradation of cohesin subunits in telophase arrested cells using *cdc15-as* allele. Cdc15-as cells and cdc15-as cells carrying MCD1-AID or MCD1-AID and SMC3-AID were synchronized in late anaphase (cdc15-as) and treated with IAA for 1 hour to deplete Mcd1-AID and Smc3-AID. Samples were taken for HiC and for immunoblotting to follow the degradation of Mcd1 and Smc3. **b**, Log_2_-ratio of contact maps between G1 (alpha factor) arrested cells and late telophase arrested cells (*cdc15-as*) (left), Log_2_-ratio of contact maps between *cdc15-as* arrested cells and *cdc15-as* arrested cells depleted in Mcd1 (right, top-half of the matrix) or Mcd1 and Smc3 (right, bottom-half of the matrix). x and y axis represent the 16 chromosomes of the yeast genome depicted on top of the matrix. Blue to red colour scales represent the enrichments in contacts in one sample respect to the other (bin=50kb). Each condition has 2 biological replicas. **c, d**, Pile-ups of contact maps of the 100 kb peri-centromeric regions in *cis c* or in trans *d* for cells synchronized in *cdc15-as* (left), cells synchronized in *cdc15as* and Mcd1 depleted (middle) cells synchronized in *cdc15as* and Mcd1 and Smc3 depleted (right). Log2-ratio of contact maps between *cdc15as* and *cdc15as* Mcd1 depleted (far right, top-half of the matrix) and *cdc15as* and *cdc15as* Mcd1 Smc3 depleted (far right, bottom-half of the matrix).

Cohesin also contributes to the organisation of the ribosomal gene array (rDNA) on chromosome XII during metaphase (Lavoie et al., 2002). We therefore tested whether inactivation of cohesin in telophase arrested cells had any effect on rDNA structure. To this aim we used an analogue-sensitive allele of *cdc15-as* and the temperature sensitive allele of cohesin’s kleisin *mcd1-73*. We expressed the nucleolar marker Net1 fused to GFP (*NET1*-yeGFP) in these cells to evaluate rDNA structure. Inactivation of cohesin in telophase arrests caused decondensation of rDNA signals (Fig. 8a). Next, we looked at chromosome XII HiC contact maps of cells arrested in Cdc15 and compared it to cells arrested in Cdc15 with either Mcd1 or Mcd1 and Smc3 degraded (Fig. 8b). These results showed a decrease in isolation of the post-rDNA region of chromosome XII and a decrease of contacts between *CEN12* and the rDNA further supporting an active role for cohesin in the organisation of specific genome regions in telophase arrested cells.

**Figure 8.**
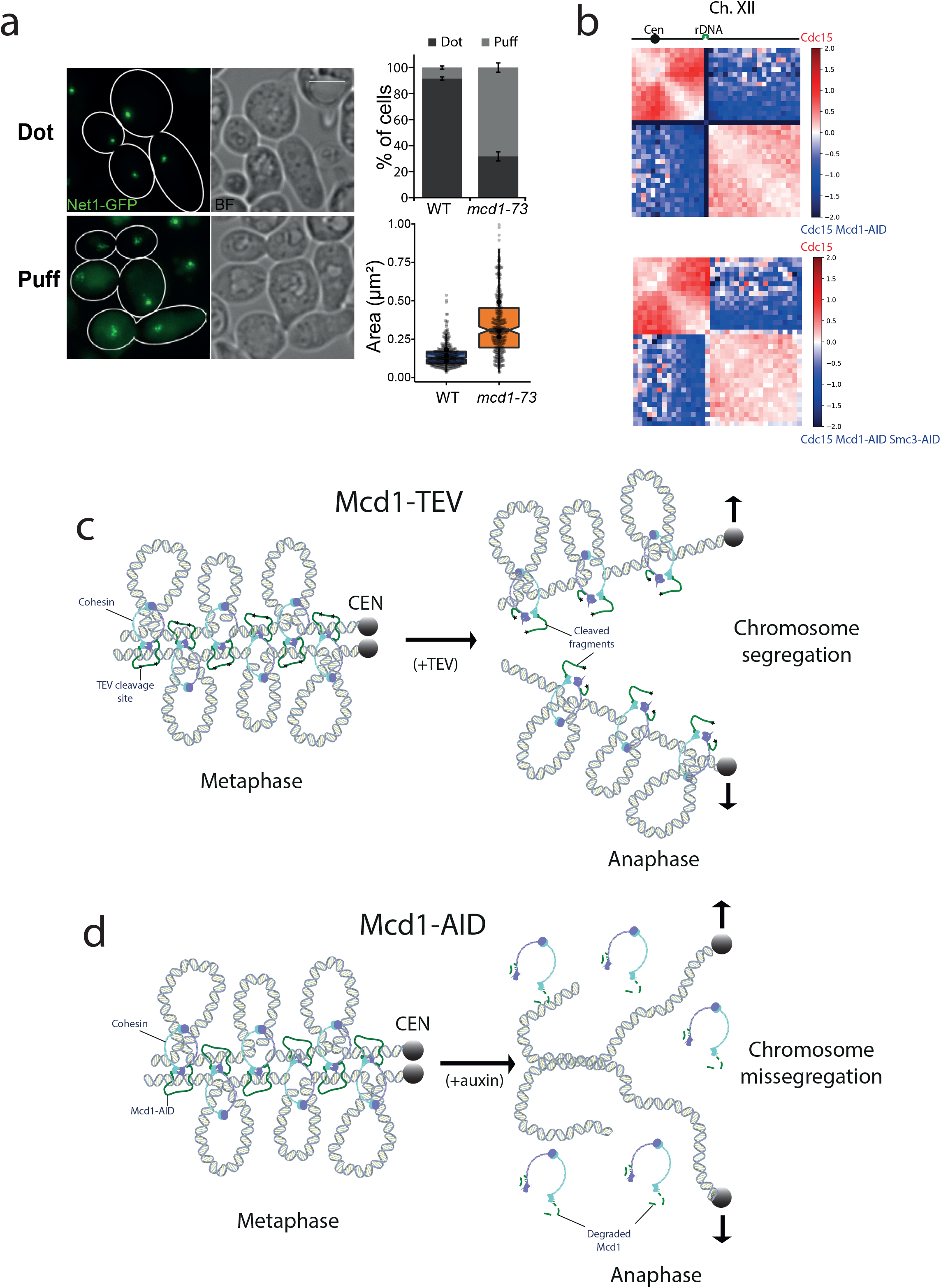
Cohesin has a structural role after metaphase. **a**, Cells containing *cdc15-as* and *NET1*-yeGFP with either MCD1 WT or the temperature sensitive allele *mcd1-73* were arrested in telophase (*cdc15-as*) at 25°C and then the temperature was shifted to 37°C for 30 minutes. Cells were then imaged under the microscope. Representative images of the experiment are shown. White scale bars represent 5 μm (left). Quantification of the rDNA morphology was scored (upper right). Net1-GFP area marking the rDNA was quantified (lower right). At least 100 cells were quantified for each condition and each condition has 3 biological replicas **b**, Log2-ratio of chromosome XII contact maps between Cdc15 and Cdc15 Mcd1-AID (top) or Cdc15 and Cdc15 Mcd1-AID Smc3-AID (bottom). **c**, Diagram showing a potential model explaining how TEV-cleaved cohesin could maintain the looped organisation of segregating chromatids. TEV cleavage of Mcd1 could be retained in one of the two segregating chromatids maintaining some of the structural functions. In this scenario sister chromatid cohesion would be lost but the loop organisation of individual chromatids would be partially maintained, thus facilitating segregation. **d**, Diagram showing a potential model explaining how degradation of cohesin subunit Mcd1 could lead to catastrophic segregation. Mcd1 degradation leads to the destabilisation of cohesin on chromatin. In the absence of cohesin, though cohesion is dissolved (allowing genome separation), the loss of structure on separated chromatids would prevent their segregation and cells would exhibit the anaphase bridges phenotypes observed.

## Discussion

The role of cohesin complexes in supporting mitotic chromosome architecture in yeast has long been established (Guacci et al., 1997; Michaelis et al., 1997), however, while their prominent role in mediating sister chromatid cohesion has been intensively studied, their contribution to chromosome folding has only recently become topical with the development of Hi-C techniques (Dekker et al., 2013; Schmitt et al., 2016). Here we set out to explore whether cohesin play a role in chromosome structuration after metaphase.

Cohesin is loaded onto chromosomes by the loader complex Scc2/4 (Ciosk et al., 2000) during G_1_ and becomes cohesive during DNA replication (Uhlmann and Nasmyth, 1998). The cohesive state requires the acetylation of cohesin Smc3 by the Eco1 acetyl-transferase to make cohesin refractory to an inhibitory “anti-establishment” activity dependent on Wapl (Rolef Ben-Shahar et al., 2008; Unal et al., 2008). In yeast, cleavage of one of cohesin subunits, Mcd1/Scc1, by the protease separase (Esp1) prompts anaphase segregation (Uhlmann et al., 1999; Uhlmann et al., 2000). Separase function is highly regulated to prevent its premature activation before every single chromosome has been aligned on the mitotic spindle. Firstly, separase activity is blocked by a bound inhibitor named securin (Pds1) (Ciosk et al., 2000), which is destroyed by ubiquitin-mediated proteolysis once cells are ready to enter anaphase (Cohen-Fix et al., 1996). Secondly, separase-mediated cleavage of Mcd1 is primed by Polo-kinase (Cdc5) dependent phosphorylation of Mcd1 at serines 175 (S175) and 268 (S268), at the cleavage site (Alexandru et al., 2001). This double regulation ensures that cohesion is not destroyed before it should be. At the anaphase onset, Mcd1 cleavage by separase is thought to terminate the function of cohesin on yeast chromosomes (Nasmyth, 2001). Interestingly, cohesin’s subunits have been shown to be chromatin-bound during anaphase/telophase (Renshaw et al., 2010; Tanaka et al., 1999), however the functional contribution of this population has not been studied.

In contrast to yeast, sister chromatid cohesion in mammals is maintained throughout G2. At the onset of mitosis, loosening of arm cohesion accompanies the compaction of each chromatid along its longitudinal axis (Waizenegger et al., 2000). Mammalian cohesin organizes the genome into loops during interphase (Rao et al., 2017) but dissociates from chromosomes in prophase (Sumara et al., 2000), and it is only found at centromeres by the time cells reach metaphase (Sumara et al., 2000). The role of organising loops on mammalian metaphase chromosomes is taken over by condensin I and II (CI and CII) complexes (Gibcus et al., 2018). During chromosome segregation, CI and CII are believed to maintain the structure of separating chromatids to prevent a “cut” phenotype, which occurs when insufficiently condensed chromatids are trapped by the cytokinetic furrow. In yeast, there is a single condensin complex that does not play a role in the overall organisation of chromosome loops during mitosis (Lazar-Stefanita et al., 2017; Schalbetter et al., 2017), instead it has a very defined function organising the ribosomal gene array on chromosome XII (Lavoie et al., 2002). Since yeast condensin does not contribute to the structural organisation of chromosome arms and cohesin’s function is thought to stop at the anaphase onset, it was unclear how separating chromatids maintain the looped organisation necessary for chromosome compaction and faithful segregation.

Here, we tested the belief that cohesin has no roles after metaphase. Our results demonstrate that, in contrast to the current view, degrading Mcd1 during anaphase leads to catastrophic segregation where many cells arrest with unseparated nuclear masses (Fig.1a-d). This result prompted us to further investigate (i) whether the structure of segregating chromatids is disrupted when Mcd1 is degraded and (ii) whether cleavage of cohesin subunit Mcd1 causes a similar phenotype to its degradation. Hi-C analysis during segregation demonstrated that Mcd1 degradation causes dramatic defects in the 10-30kb range of *cis* interactions on chromatids, which correspond to cohesin-dependent loop range (Fig. 5c). This result confirms that loops are lost when cohesin is not present on segregating chromatids and offers an explanation for the catastrophic segregation observed under these conditions (Fig. 1c-d).

To study the effect of cohesin cleavage on the efficiency and structure of segregating chromatids we used the TEV protease system previously described (Uhlmann et al., 2000). Our analysis revealed that unlike Mcd1 degradation, TEV-induced cleavage does not prevent bulk nuclear separation (Fig. 2 & 4), although it causes some segregation errors (Fig. 4c). Importantly, we could detect the maintenance of *cis* contacts in 10-30kb range on chromosomes with TEV-cleaved Mcd1 (Fig. 5c) demonstrating that chromatid loops are not fully disrupted and explaining why bulk segregation is achieved. An important difference between cohesive cohesin and loop extruding complexes lies in the way that these complexes interact with the DNA substrate. While cohesive cohesin topologically embraces DNA (Srinivasan et al., 2018), loop extruding complexes rely on non-topological interactions (Davidson et al., 2019). Therefore, it is likely that cohesin complexes cleaved by TEV are still able to interact non-topologically with the DNA substrate (Fig. 8c) and/or can maintain extruded loops. This would explain why TEV-cleavage has only a modest effect on the structural role of cohesin, while disrupting the cohesive role (that depends on topological association) (Fig. 8c). In contrast, auxin-mediated degradation would abolish both topological and non-topological interactions (Fig. 8d) which would result in the loss of cohesion but also structure (and compaction) thus generating a situation where unorganised chromatids would fail to efficiently segregate (Fig. 8d).

Based on evidence that suggested a role for cohesin during segregation we sought to investigate whether a small pool of cohesin was retained on chromosomes in telophase. Interestingly, we detected binding to centromeres in this late mitotic stage (Fig. 6a and c). Importantly, depletion of cohesin in telophase revealed that bound cohesin during this late mitotic stage is still influencing the structural organisation of pericentromeric regions, as well as promoting interactions between *CEN*-regions of different chromosomes (Fig. 7b-d). Further to this role at centromeric regions, we observed condensation defects at the ribosomal gene array on chromosome XII when we inactivated cohesin function in telophase-arrested cells (Fig. 8a-b) demonstrating that the complex also plays a role in the organisation of this genomic site during late mitosis.

Currently, the roles of yeast cohesin during mitosis are thought to include; (i) the pairing of sister chromatids (Guacci et al., 1997; Michaelis et al., 1997), (ii) the bipolar organisation of chromatids as they attach to the mitotic spindles (Tanaka et al., 2000) and (iii) the organisation of metaphase chromosomes into looped domains (Lazar-Stefanita et al., 2017; Schalbetter et al., 2017). Recent work has shown that cohesin is necessary for the maintenance of the structure of the rDNA during metaphase and that this role is executed through the regulation of condensin localisation (Lamothe et al., 2020). Importantly, all the above roles occur in metaphase cells and the cleavage by separase at the anaphase onset has generated the perception that cohesin’s roles are over at the beginning of anaphase. Collectively our data demonstrates that yeast cohesin plays important roles during segregation by maintaining the looped organisation of segregating chromatids and supporting centromere organisation. Therefore, these findings extend the repertoire of cohesin jobs on chromosomes and demonstrates that this complex has important post-metaphase functions critical for ensuring faithful genome segregation.

## Methods

### Yeast strain

Yeast strains used in this study are listed in Table 1. Epitope tagging of genes were carried out as described in (Janke et al., 2004).

|  |  |  |  |
| --- | --- | --- | --- |
| CCG14820 | CCG4000; *MET3-3HA-CDC20:TRP1; ADH1-OsTIR1::URA; MCD1:AID-9Myc:Hph; TetR-GFP::ADE2; TetO:469Kb ChrV:HIS3* | This study | Fig 1a, 1b, 1c |
| CCG14881 | CCG4000; *MET3-3HA-CDC20:TRP1; ADH1-OsTIR1::URA; MCD1:AID-9Myc:Hph; TetR-GFP::ADE2; TetO:558Kb ChrV:HIS3* | This study | Fig 1d |
| CCG14629 | CCG4000; *MET3-3HA-CDC20:TRP1; ADH1-OsTIR1::URA; MCD1:AID-9Myc:Hph; TetR-GFP::ADE2; TetO:448Kb ChrIV:HIS3* | This study | Fig Supp 1a |
| CCG14635 | CCG4000; *MET3-3HA-CDC20:TRP1; ADH1-OsTIR1::URA; MCD1:AID-9Myc:hph; TetR-GFP::ADE2; TetO:1513Kb ChrIV:HIS3* | This study | Fig Supp 1b |
| K9027 | *MATα; MCD1Δ:HIS3; MCD1TEV268::LEU2; GAL-NLS-mcy9-TEVprotease-NLS2::TRP1 (10-fold integrant by southern); ura3::3xURA3 tetO112; his3::HIS3 tetR-GFP; MET3-HA3-CDC20::TRP1; ade2-1 can1-100 GAL psi+* | Uhlmann (2000) | Fig 2a, 2b and 2c |
| CCG12955 | CCG12681; SMC3-V5:HPH | This study | Fig 3 |
| CCG12356 | CCG4000; *MET3-3HA-CDC20:TRP1, FLAG-MCD1-268TEVF-6HA:LEU2, GAL-TEV-9Myc::URA3; tetR-GFP::ADE2* | This study | Fig 4a, 4b |
| CCG12351 | CCG4000; *MET3-3HA-CDC20:TRP1, FLAG-MCD1-268TEVG-6HA:LEU2, GAL1-TEV-9Myc::URA3; tetR-GFP::ADE2* | This study | Fig 4a, 4b, 5b, 5c<br>Supp. 4 & 5 |
| CCG14783 | CCG12351; *TetO:469Kb ChrV:HIS3* | This study | Fig 4c |
| CCG14784 | CCG12356; *TetO:469Kb ChrV:HIS3* | This study | Fig 4c |
| CCG14882 | CCG14635; SMC3-V5:hph | This study | Fig 5a |
| CCG12925 | CCG12682; SMC3-V5:HPH | This study | Fig 5a |
| CCG13574 | CCG4000, *MET3-3HA-CDC20:TRP1*; pADH1-*OsTIR1::URA*; *TetR-GFP::ADE2* | This study | Fig 5b, 5c<br>Supp. 4 & 5 |
| CCG14635 | CCG4000; *MET3-3HA-CDC20:TRP1*; *ADH1-OsTIR1::URA*; *MCD1:AID-9Myc:hph*; *TetR-GFP::ADE2*; *TetO:1513Kb ChrIV:HIS3* | This study | Fig 5b, 5c<br>Supp. 4 & 5 |
| CCG14880 | CCG4000; *ADH1-OsTIR1::LEU*; *CDC15:AID-9Myc:Hph* | This study | Fig 6a |
| CCG14909 | CCG4000; *ADH1-OsTIR1::LEU*; *CDC15:AID-9Myc:Hph* SMC3-yeGFP:Kan | This study | Fig 6a |
| CCG14910 | CCG4000; *ADH1-OsTIR1::LEU*; *CDC15:AID-9Myc:Kan*; SMC3-yeGFP:Hph; *SPC29-RedStar2:Nat* | This study | Fig 6a |
| CCG1835 | *MATa bar1:hisG ura3-1 trp1-1 leu2-3,112 his3-11 ade2-1 can1-100 GAL+ cdc15-2* | This study | Fig 6b<br>Supp. 6 & 7 |
| CCG14731 | CCG1835; *MCD1-6HA:Hyg* | This study | Fig 6b<br>Supp. 6 & 7 |
| CCG14758 | CCG4000; *SMC1-6HA:Nat*; pADH1-*OsTIR1::URA* | Garcia-Luis J.<br>*NSMB*<br>(2019). | Fig 6b<br>Supp. 6 & 7 |
| CCG14821 | CCG4000 pADH1-*OsTIR1::LEU*; *cdc15as1:KanMX* | This study | Fig 7a, 7b, 7c<br>Supp. 8 |
| CCG14822 | CCG14821 *MCD1:AID-9Myc:Hph* | This study | Fig 7a, 7b, 7c<br>Supp. 8 |
| CCG14823 | CCG14822 *SMC3:AID-9Myc:Hph* | This study | Fig 7a, 7b, 7c<br>Supp. 8 |

### Media, culture conditions and DNA constructs

To arrest the cells in G1, α-factor was added to exponentially growing MATa cultures (OD600=0.5) to a final concentration of 3 x 10^-8^ M for 3 hours at 25°C. To arrest cells in G2/M, Nocodazole (1.5 mg/mL stock in DMSO 100%) was added to cultures with OD600=0.5 to a final concentration of 0.015 mg/mL for 2.5 hours. To arrest cells in metaphase (Cdc20), cells carrying the *CDC20* gene under methionine repressible promoter MET3 (*MET3-CDC20*) were grown overnight in minimal media lacking methionine. The day after cells were arrested by washing the cells and resuspending them in rich media containing 5mM methionine for 3 hours. To arrest cells in telophase, Cdc15 was tagged with an auxin degron (*CDC15-AID*), and IAA was added to the culture at a final concentration of 3mM or 1mM when growing in minimal media. To arrest the cells in late telophase using the cdc15-as allele, cells released from an alpha factor arrest were treated with 10μM of the ATP analogue 1-NM-PP1 for 1 hour 45 minutes. Cultures were monitored by microscopy until ≥90% of cells were arrested. To release cells from G1, the culture was spun (4,000 r.p.m, 1 min) and washed in YPD 3 times. The pellet was then resuspended in YPD containing 0.1 mg/mL pronase. To release cells from Nocodazole, the culture was spun (4,000 r.p.m, 1 min) and washed in YPD containing 1% DMSO 5 times. The pellet was then resuspended in YPD. To degrade proteins tagged with AID epitope, a stock of IAA of 0.6M in ethanol 100% was used.

### Microscopy and statistics

To monitor cell cycle progression and chromosome segregation an epifluorescence OLYMPUS IX70 microscope was used fitted with a Lumecor Spectra LED light source, a Hamamatsu Orca Flash 4.0 V2 camera and a 100X/1.35 lens. 1 mL of cell culture was taken from each time point and mixed with glycerol (20% final concentration) to preserve *TetO*/TetR signal after being frozen at −80°C. For visualization, cells were centrifuged at 3,000 r.p.m. for 2 minute and ~ 1 μL of the pellet was mixed with ~1 μL of DAPI solution (DAPI 4 μg/mL Triton 1 %) on the microscope slide. For each field 20 z-focal planes images were captured (0.3 μm depth between each consecutive image). Images were analyzed with Fiji (Schindelin et al., 2012). To quantify the distance between the GFP dots a Fiji macro was developed to automatically compute the weighted centroid of the dots and measure the three-dimensional distance between them. To visualize Net1-yeGFP or Smc3-yeGFP, cell were imaged fresh in a DELTAVISION Elite fluorescence microscope fitted with a Lumecor Spectra LED light source, a Photometrics Coolsnap HQ camera and a 100x/1.4 lens. Smc3-yeGFP signal intensity was calculated as the ratio of signal at the centromeres and the signal of the same area in the nucleus on Z-projection of images taken every 0.2 μm in 6 μm. In the box plot, centre lines represent the medians; box limits indicate the 25th and 75th percentile and notches give roughly 95% confidence intervals for each median. Whisker extends from the hinge to the largest value no further than 1.5 * IQR. Dots outside the whiskers represent outliers. When grey transparent dots are shown they represent individual values of each cell. Histograms to show rDNA morphology represent the average of 3 biological replicas and error bars the standard deviation.

### Western blot

Protein extraction was done by lysing the cells in a FastPrep FP120 (BIO101) machine with 20% TCA and glass beads. 3 repetitions of a 20 seconds cycle, power setting 5.5. Proteins were precipitated with TCA 7.5% and centrifuging at 15000rpm for 10 minutes at 4 °C. Then the pellet was resuspended in Laemmli buffer 1.5X. Western blots were resolved in 7.5% SDS-PAGE gels. Proteins were transferred to polyvinylidene fluoride (PVDF) membranes using the TE70X semidry blotter system (Hoefer). The antibodies used were anti-HA (Roche, 3F10), anti-Myc (Roche, 9E10), anti-PGK1 (Thermo Scientific, 459250) anti-V5 (Abcam, ab9116) and anti-FLAG (Invitrogen MA1-142). Blots were incubated with the ECL Prime Western blotting detection reagent (GE Healthcare). Blots were developed by exposure to high-performance chemiluminescence films (Amersham Hyperfilm ECL, GE Healthcare) or in an ImageQuant LAS 4000 mini machine (GE Healthcare).

### Chromatin Immunoprecipitation

For ChIP analysis, cells were grown to OD600 = 0.5 and arrested at the required cell cycle stage. A total of 100 OD600 units of S. cerevisiae were collected. Cells were fixed for 15 min at 25 °C and quenched with glycine (final concentration 125 mM) for 7 min before cells were harvested by centrifugation at 4,000 r.p.m. for 1 min. The cell pellets were washed in PBS and transferred to a screw cap tube and frozen on dry ice. The pellets were stored at −80 °C. Pellets were resuspended in 300 μl of IP buffer (150 mM NaCl, 50 mM Tris-HCl (pH 7.5), 5 mM EDTA, NP-40 (0.05% vol/vol), Triton X-100 (1%vol/vol)) containing PMSF (final concentration 1 mM) and complete protease inhibitor cocktail (without EDTA, from Roche). A 500 μl volume of glass beads was added to the tubes. Cells were broken in a FastPrep FP120 cell disruptor (BIO101) by three repetitions of a 20 s cycle at power setting 5.5. The cell lysate was transferred to a new tube and 100 μl volume of IP buffer containing PMSF and protease inhibitors was added. The cell lysate was spun down for 10 min at 15,000 r.p.m at 4 °C. This pellet was resuspended in 1 ml of IP buffer containing PMSF and protease inhibitors, and sonicated for 30 (30 s on, 30 s off) at high power at 4 °C in a Diagenode Bioruptor pico. After sonication samples were spun down for 10 min at 15,000 r.p.m. the supernatant was taken. A 200 μl volume of the sonicated chromatin was taken as ‘input’ and 400 μl was incubated with 40 μg of anti-V5 antibody (anti-V5 Abcam, ab9116) in a sonicator at low power for 30 min (30 s on, 30 s off). The ‘input’ DNA was precipitated with 0.3 M sodium acetate and 2.5 volumes of cold ethanol and spun down at 15,000 r.p.m. for 30 min, then the supernatant was removed. The pellet was washed with 70% ethanol and air-dried. After antibody binding, the IP sample was spun down at 13,000 r.p.m for 5 min and the supernatant was added to 60 μl of Dynabeads protein G (Invitrogen), previously equilibrated with IP buffer. The samples were then incubated for 2 h at 4 °C in a rotating wheel and washed five times with IP buffer using a magnetic separator rack. Finally, ‘input’ samples and IP samples were resuspended in de-crosslinking buffer (TE 1×, 1% SDS, 10 μg ml-1 RNase A, 1 mg ml-1 proteinase K) and incubated at 65 °C overnight. Samples were purified using a ChIP DNA Clean & Concentrator kit (Zymoresearch) according to the manufacturer’s instructions. Primers used for ChIP-qPCR are described in the table below. Calibrated ChIP-seq were done as described in (Garcia-Luis et al., 2019).

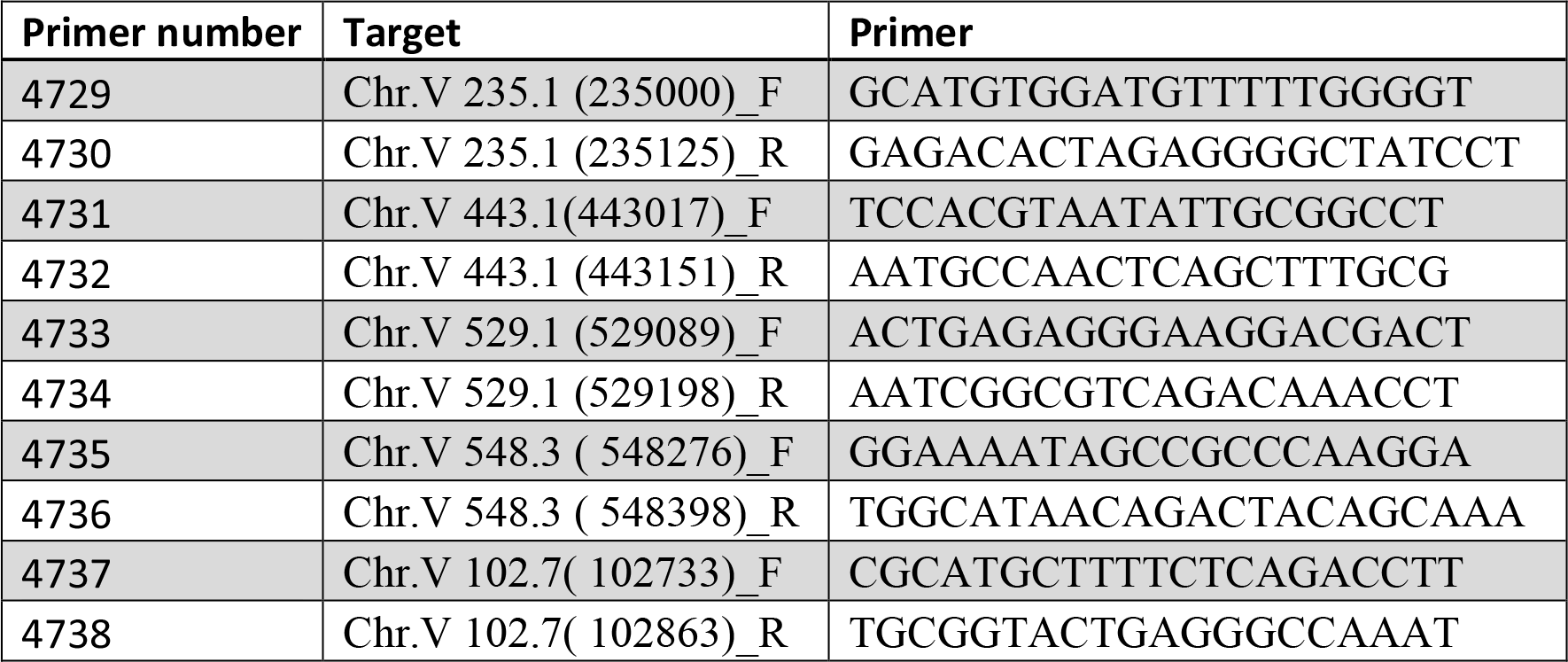
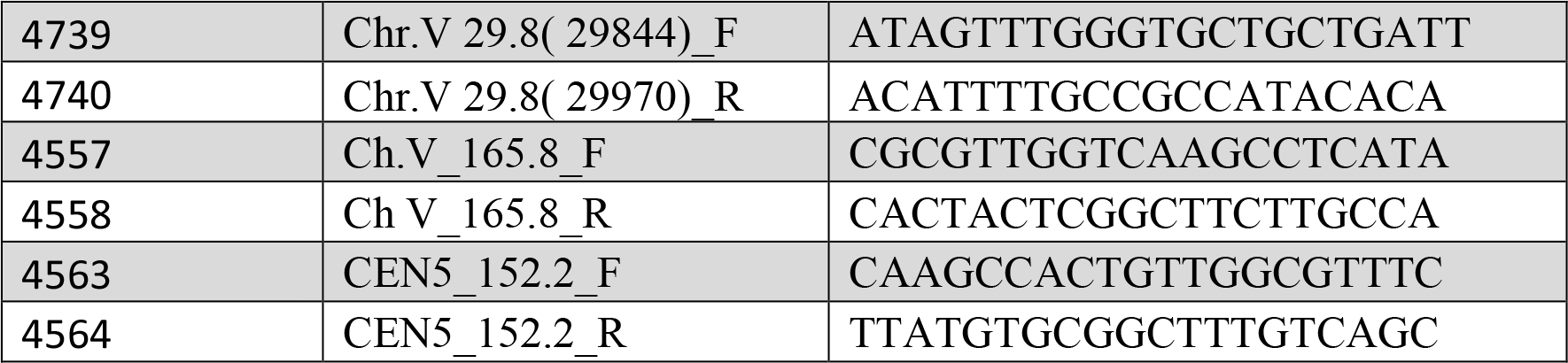

### Protein co-immunoprecipitation

120 OD_600_ of asynchronous cells were OD_600_ = 1 were collected and washed in cold water and resuspended in 200 μl of ice-cold buffer A (50 mM HEPES, 150 mM KCl, 1.5 mM MgCl2, 0.5 mM DTT and 0.5% Triton X-100 (pH 7.5) supplemented with complete protease inhibitor cocktail tablets (Roche)). 500 ml of glass beads (425-600 μm) were added and cells lysed in a FastPrep FP120 cell disruptor (BIO101) by three repetitions of a 20 s cycle at power setting 5.5. Extracts were maintained on ice for 2 min after each cycle. Cell extracts were centrifuged for 10 minutes at 12000 r.p.m at 4C and the supernatant incubated with protein G Dynabeads (Invitrogen) bound to anti-Myc antibody (Roche, 9E10) for 2 h at 4 °C. Finally, beads were washed five times in washing buffer (10 mM Tris-Cl pH 7.5, 150 mM NaCl, 0.5% Triton) and unbound from the antibody by incubating at 37°C for 4 min in SR buffer (2% SDS, 0.125 M Tris-Cl pH 6.8). Immunoprecipitated proteins were mixed with SS buffer (5% saccharose, 0.0125% bromophenol blue) and run in an SDS–PAGE gel.

### Hi-C libraries

Cells were fixed with 3% formaldehyde (REF) as detailed in (Dauban et al., 2020). Formaldehyde was quenched with 300 mM of glycine at room temperature for 20 min. Hi-C experiments were performed with a Hi-C kit (Arima Genomics) involving a double DpnII + HinfI restriction digestion. Preparation of the samples for paired-end sequencing on an Illumina NextSeq500 (2×35 bp) was done with Collibri™ ES DNA Library Prep Kit for Illumina Systems (Thermo Fisher, A38605024).

### Generation and normalization of contact maps

Alignment of the reads and processing of the contact data was done with Hicstuff using the S288C reference genome (Matthey-Doret et al., 2020). Hicstuff pipeline was launched with the following parameters: filter out spurious 3C events, filter out PCR duplicates based on read positions. The “view” mode of Hicstuff SCN function was used to generate normalized contact maps as described in (Cournac et al., 2012). Contact maps were binned at 50kb for the whole genome or 1kb for single chromosomes, and 30 kb for single chromosome ratio map.

### Contact probability as a function of the genomic distance P(s)

Genome-wide contact probability as a function of genomic distance Pc(s) and its derivative were computed using the “distance law” function of Hicstuff with default parameters, averaging the contact data of entire chromosome arms (Matthey-Doret et al., 2020). For pre-rDNA region, contact probability as a function of genomic distance Pc(s) and its derivative were computed after alignment of the reads against the 290kb long pre-and post-rDNA region, as if they were individual chromosomes.

### Aggregated of *cis* and *trans* centromere contacts

*Cis*-centromere pile-up contact maps are the result of averaged 205 kb windows with a bin of 5 kb generated with Chromosight in quantify mode (option: -- pattern border, -- perc-zero=100) (Matthey-Doret et al., 2020) centered on the 16 centromere position, and divided by the mean for each pixel of its diagonal. *Trans*-centromere pile-up were similarly generated by Chomosight, but with the option --inter, and centered on centromere intersections.

## Acknowledgements

We thank members of our laboratories for discussion and critical reading of the manuscript. We thank Helle D Ulrich for kindly sharing the plasmids for auxin-inducible degron tagging. We also thank MRC-LMS microscopy facility for help with the microscope set-up and especially Chad Whilding for developing the Fiji macro for automated distance quantification of GFP dots.

## Author contributions

J.G.L. performed all experiments except for HiC analyses. H.B. and A.T performed HiC experiments and analysis. H.B analysed HiC data. L.A. and J.G-L. conceived the project. L.A. wrote the manuscript. L.A. and R.K. supervised the projects and revised the manuscript.

## Data and materials availability

ChIP-seq and HiC data supporting the findings of this study have been deposited in the GEO database and are accessible through accession no. GSE183481. ChIP-seq of nocodazole arrest has been reused from the dataset with accession no. GSE118534. Any further data that support the findings of this study are available from the corresponding authors upon request.

