## Supplementary figures for "The contribution of cohesin to chromatid organisation is critical during chromosome segregation"

### Supplementary Figure 1

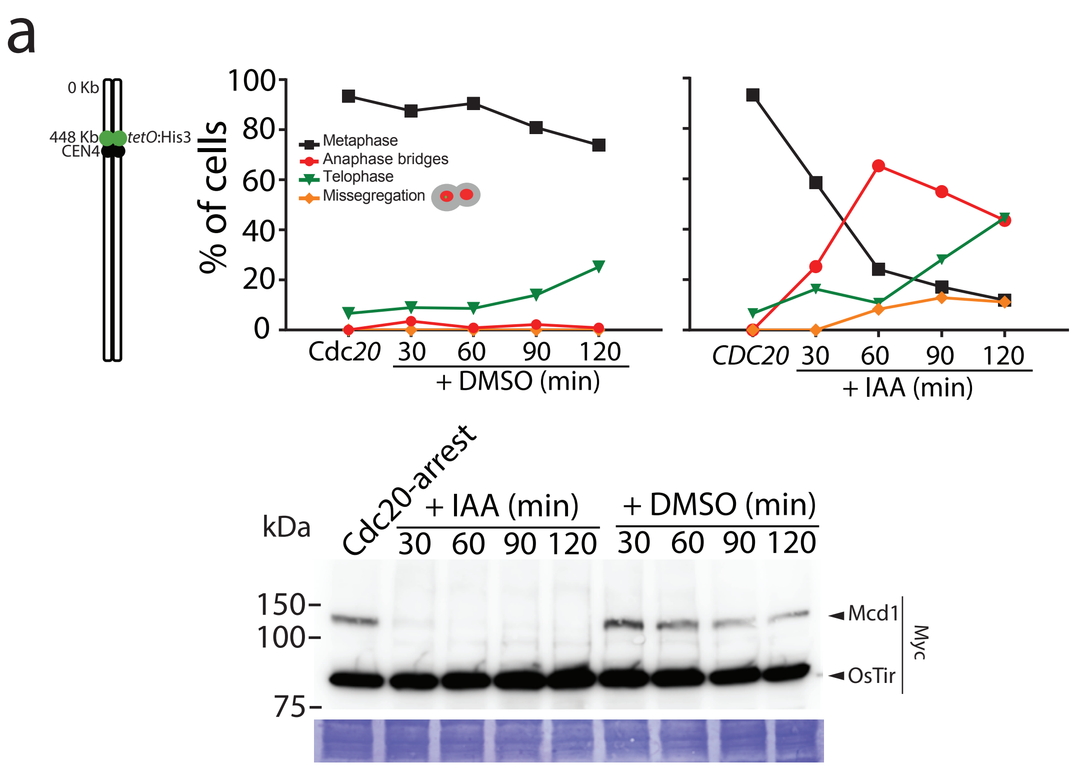


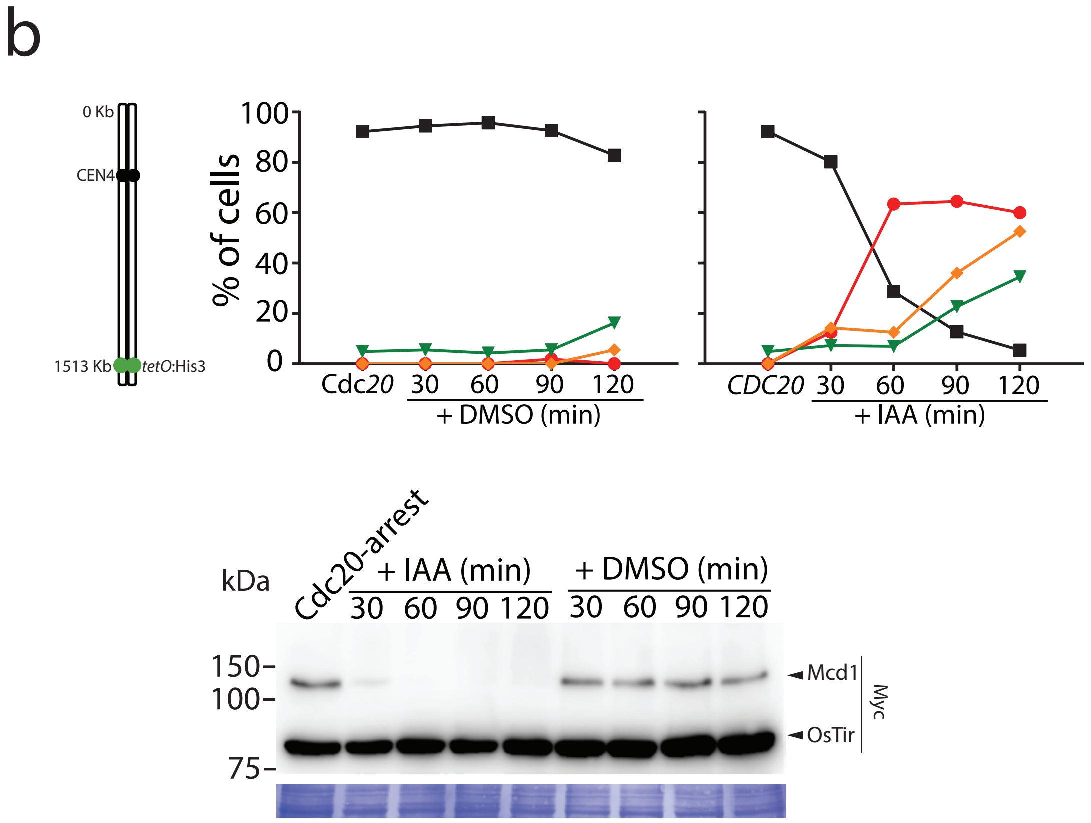

### Supplementary Figure 2

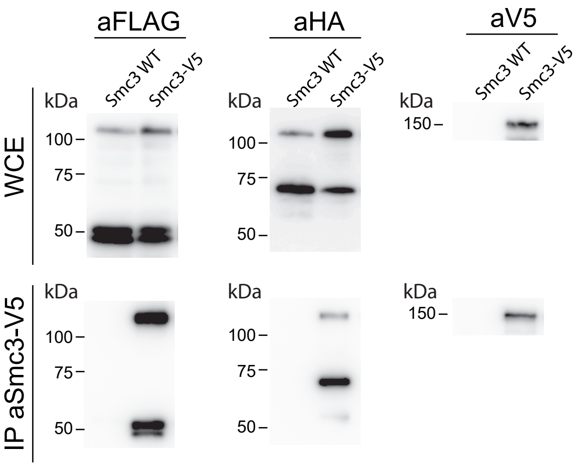

Supplementary Figure 3

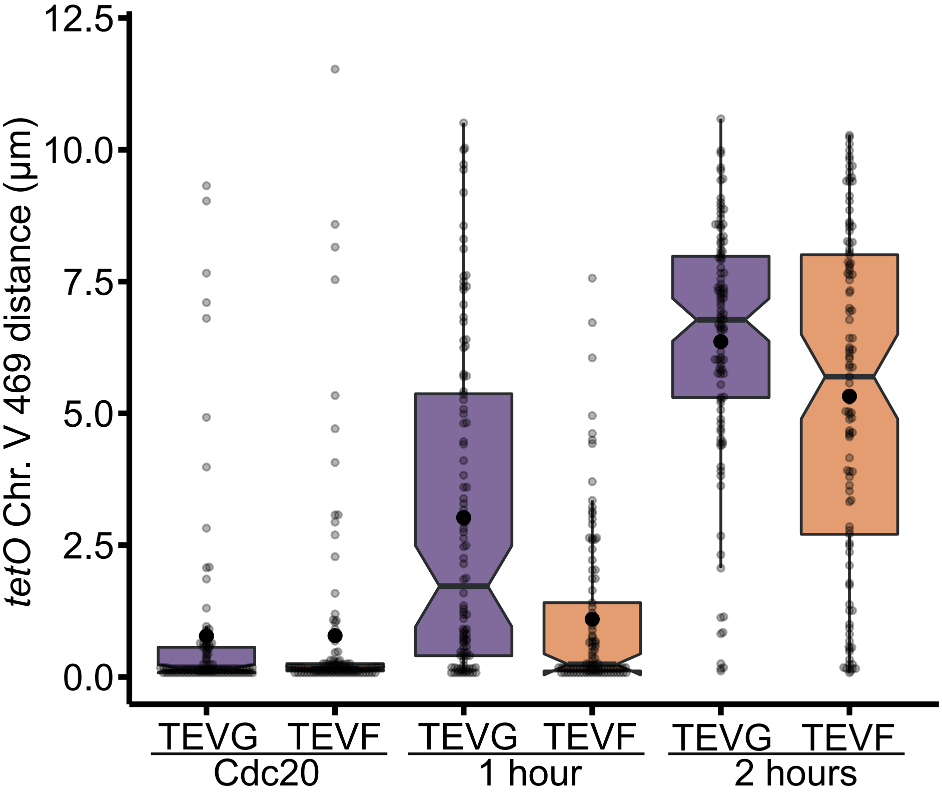

### Supplementary Figure 4

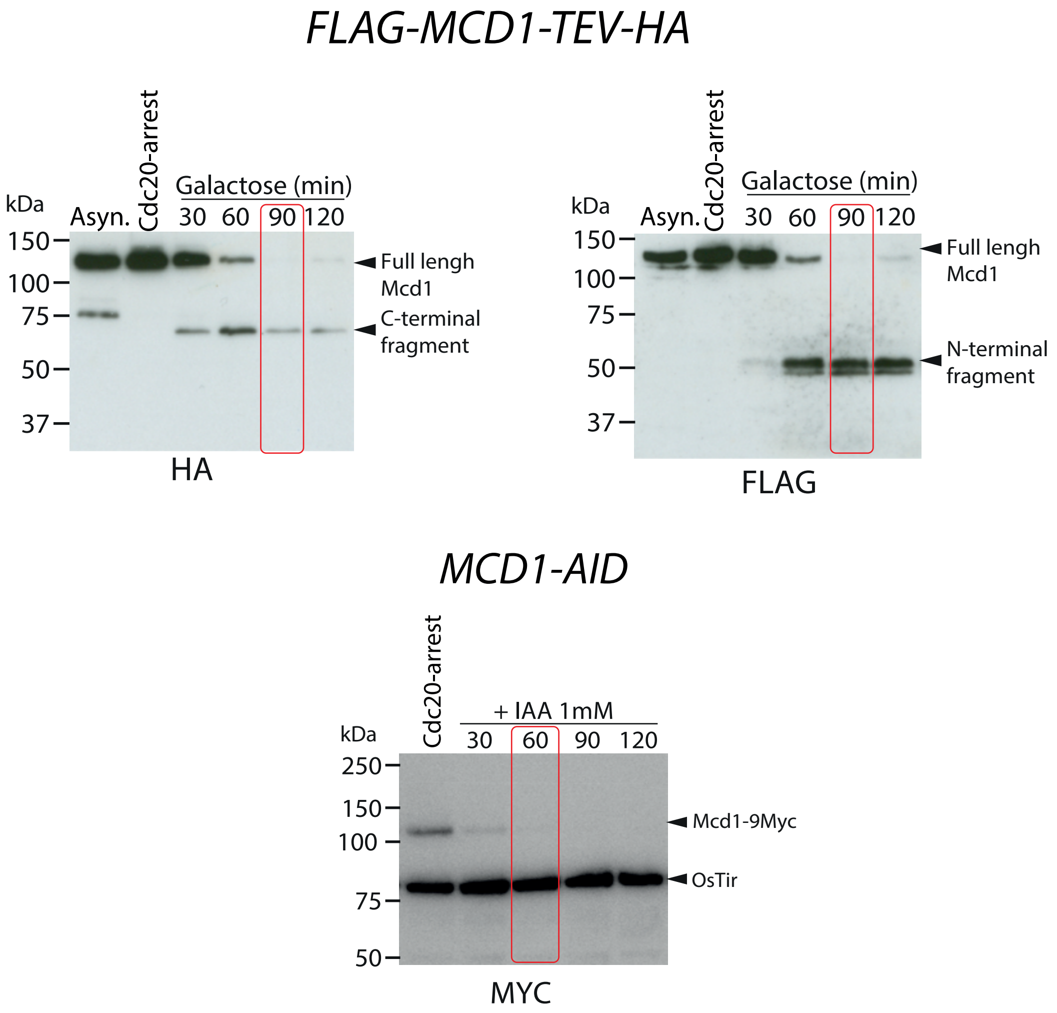


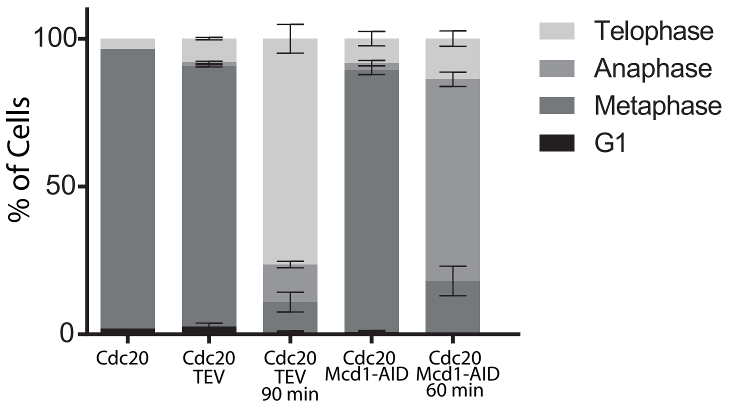


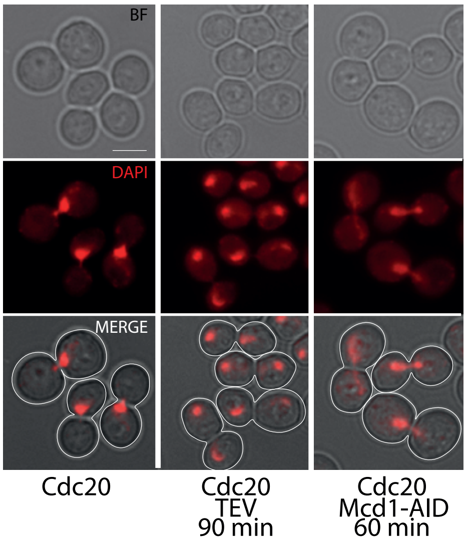

### Supplementary Figure 5

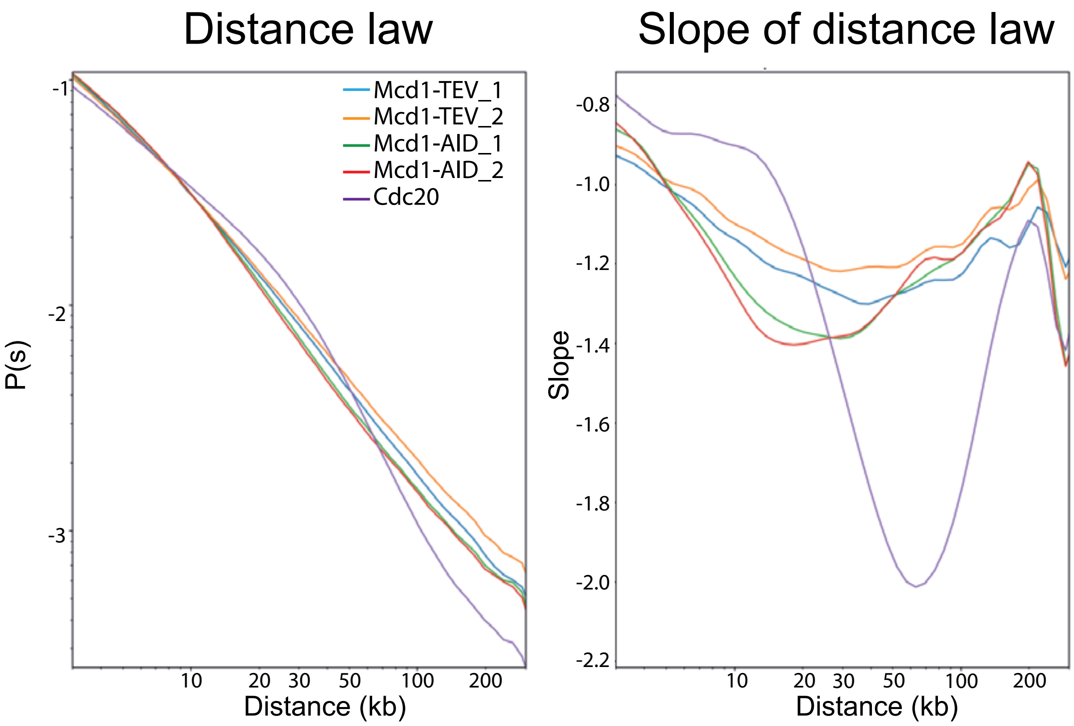

Supplementary Figure 6

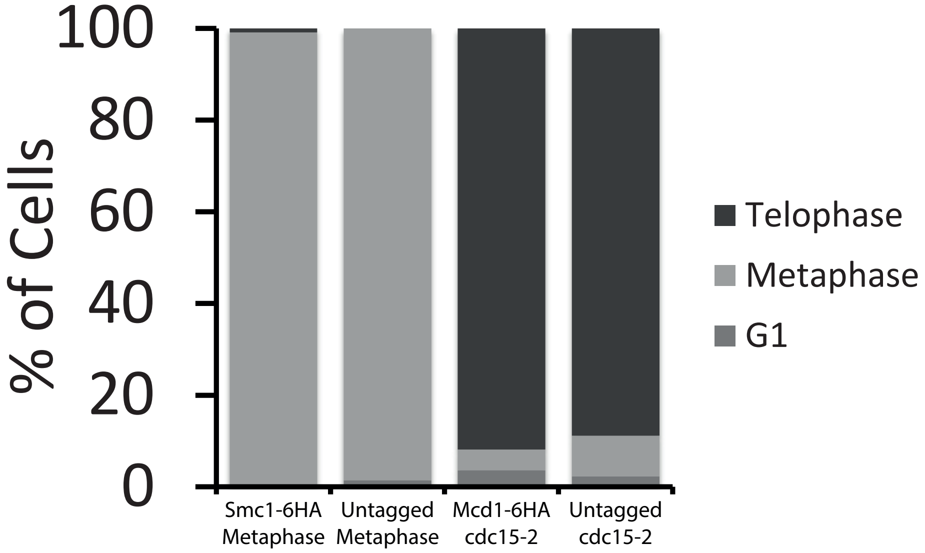

### Supplementary Figure 7

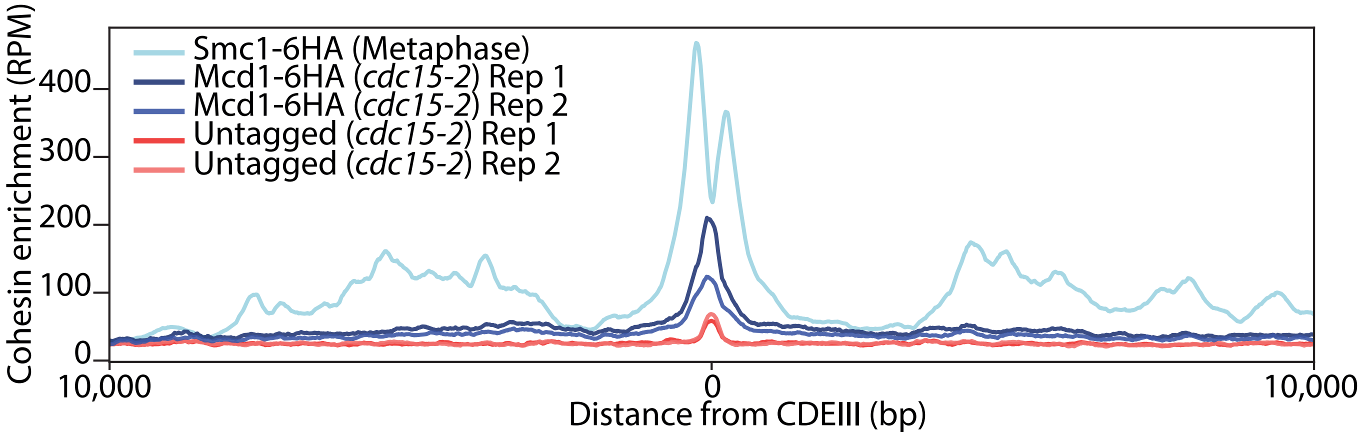

### Supplementary Figure 8

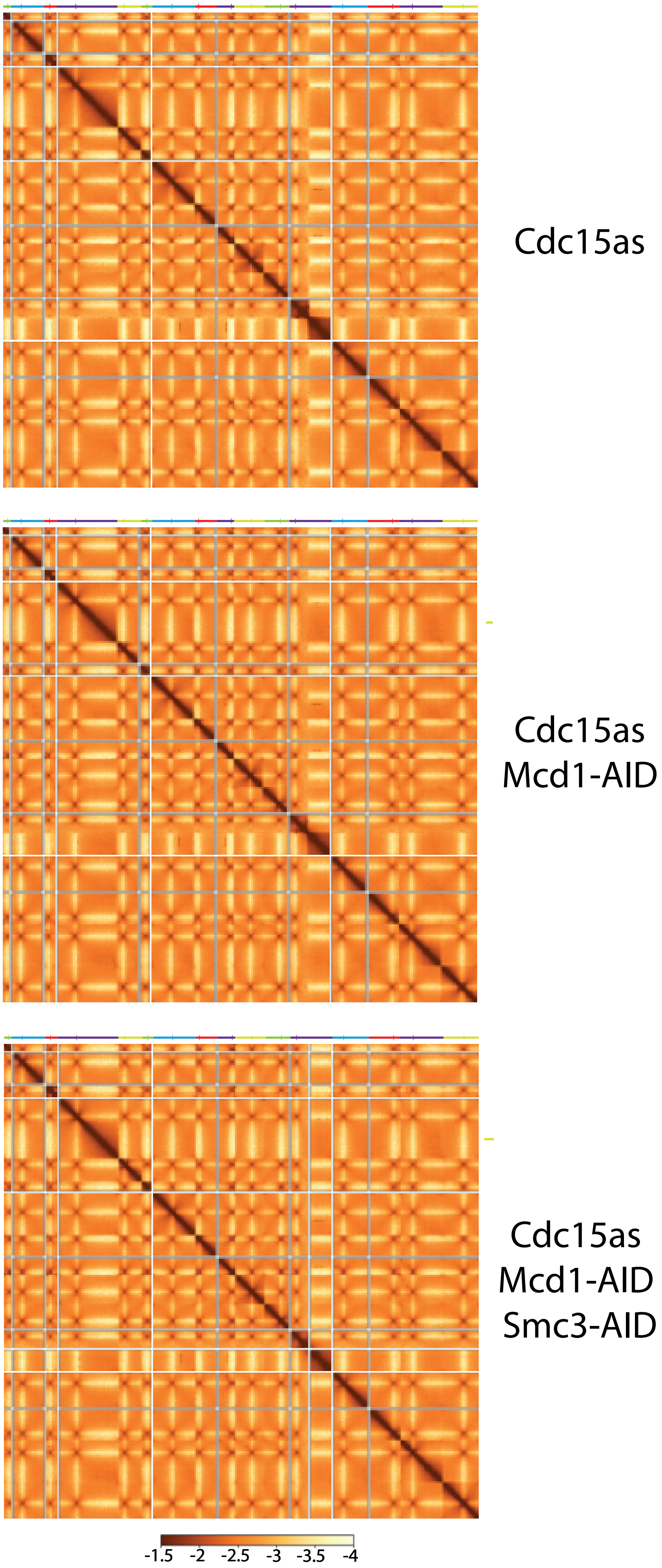
