## Supplementary figures legend for "The contribution of cohesin to chromatid organisation is critical during chromosome segregation"

**Supplementary Figure 1 (related to main Fig. 1)**

**a**, Cells with *MCD1* tagged with the auxin degron (*MCD1-AID*) and with the centromere of the chromosome IV *(tetO::448*) marked with GFP were synchronized in metaphase (Cdc20). The culture was split in two, one of them was treated with DMSO 1% and the other with 1mM of IAA to degrade Mcd1-AID. Samples were collected at the indicated timepoints and chromosome segregation analysis (top) and immunoblotting to follow Mcd1-aid (bottom) was performed. **b**, Cells treated as in a, but with the subtelomere of chromosome IV (*tetO::1513*) marked with GFP. Each time point represents the percentage of cells at the indicated cell cycle stage. At least 100 cells were quantified for each time-point.

**Supplementary Figure 2 (related to main Fig. 3)**

Cells with *MCD1* tagged in N-terminus with FLAG, C-terminus with HA, with the 268 separase cleavage position substituted by the TEV cleavage sequence ENLYFQG (TEV-G) and with *SMC3* tagged in C-terminus with V5 were synchronized in metaphase (Cdc20) and Mcd1 was cleaved in the TEV cleaving site. Samples were taken after 2 hours of induction of the TEV protease and Smc3-V5 was pulled down using anti-V5 antibody followed by anti-FLAG or HA immunoblotting to detect Mcd1 N-terminus or C-terminus cleaved fragments respectively. A negative control without V5 tag in Smc3 was included.

**Supplementary Figure 3 (related to main Fig. 4)**

Cells carrying MCD1 with the 268 separase cleaving position substituted by the TEV cleavage sequence ENLYFQF (TEV-F) or ENLYFQG (TEV-G) and with the middle of chromosome V (*tetO* 469) marked with GFP were synchronized in metaphase (Cdc20) and Mcd1 was cleaved in the TEV cleaving site. Samples were collected at the indicated timepoints and the distance between the GFP dots was quantified.

**Supplementary Figure 4 (related to main Fig. 5b-c)**

Cells containing either *MCD1 TEVG or MCD1-AID* were arrested in Cdc20 metaphase arrest and MCD1 was cleaved or degraded respectively. Samples for WB analysis were taken every 30 minutes for 2 hours followed by immunoblotting to detect Mcd1. Samples at the timepoints highlighted in red (*MCD1 TEV-G* 90 min. cleavage and *MCD1-AID* 60 minutes degradation) were taken to be processed for Hi-C (top and middle blots). The cell cycle stage of these samples was quantified (bottom left). Representative cell images of each sample are shown (bottom right).

**Supplementary Figure 5 (related to main Fig. 5c)**

Contact probability (P(s)) of cells containing either *MCD1 TEVG or MCD1-AID* synchronized in metaphase (Cdc20) and cleaved for 90 min. or degraded for 60 min. respectively. Biological replicas are shown.

**Supplementary Figure 6 (related to main Fig. 6b)**

Microscope quantification of the cell cycle stage of cells used for ChIP-seq experiments, containing SMC1-6HA arrested in metaphase (nocodazole), untagged cells arrested in metaphase (nocodazole), cells containing MCD1-6HA arrested in *cdc15-2* late anaphase and untagged cells arrested in *cdc15-2*.

**Supplementary Figure 7 (related to main Fig. 6c)**

Enrichment of cohesin around CENs measured by calibrated ChIP-seq. Smc1-6HA ChIP-seq profiles of SMC1-6HA wild-type cells synchronized in metaphase, Mcd1-6HA ChIP-seq profiles of *cdc15-2* cells synchronized in telophase and ChIP-seq profiles of untagged *cdc15-2* cells synchronized in telophase are shown. The number of reads at each base pair from the centromere CDEIII was averaged over all 16 chromosomes. The profile of each biological repetition is shown.

**Supplementary Figure 8 (related to main Fig. 7b)**

Contact maps generated from cells synchronized in *cdc15-as* and cells synchronized in *cdc15-as* containing *MCD1-AID* or *MCD1-AID* and *SMC3-AID.* Cells were synchronized in late anaphase (*cdc15-as)* and treated with IAA for 1 hour to deplete Mcd1-AID and Smc3-AID. Samples were taken for HiC and for WB. x – and y – axis represent the 16 chromosomes of the yeast genome, displayed above the maps. Brown to yellow colour scales represent high to low contact frequencies, respectively (log10).
